# Lack of effect of different pain-related manipulations on opioid self-administration, reinstatement of opioid seeking, and opioid choice in rats

**DOI:** 10.1101/2021.02.11.430808

**Authors:** David J Reiner, E Andrew Townsend, Javier Orihuel Menendez, Sarah V Applebey, Sarah M Claypool, Matthew L Banks, Yavin Shaham, S Stevens Negus

## Abstract

**Rationale and Objective:** Pain-related factors increase risk for opioid addiction, and opioid-induced pain relief may function as a negative reinforcer to increase opioid taking and seeking. However, experimental pain-related manipulations generally do not increase opioid self-administration in rodents. This discrepancy may reflect insufficient learning of pain-relief contingencies or confounding effects of pain-related behavioral impairments. Here we determined if pairing noxious stimuli with opioid self-administration would promote pain-related reinstatement of opioid seeking or increase opioid choice over food.

**Methods:** In Experiment 1, rats self-administered fentanyl in the presence or absence of repeated intraplantar capsaicin injections in distinct contexts to model context-specific exposure to cutaneous nociception. After capsaicin-free extinction in both contexts, we tested if capsaicin would reinstate fentanyl seeking. In Experiment 2, rats self-administered heroin after intraperitoneal (i.p.) lactic acid injections to model acute visceral inflammatory pain. After lactic acid-free extinction, we tested if lactic acid would reinstate heroin seeking. In Experiment 3, we tested if repeated i.p. lactic acid or intraplantar Complete Freund’s Adjuvant (CFA; to model sustained inflammatory pain) would increase fentanyl choice over food.

**Results:** In Experiments 1-2, neither capsaicin nor lactic acid reinstated opioid seeking after extinction, and lactic acid did not increase heroin-induced reinstatement. In Experiment 3, lactic acid and CFA decreased reinforcement rate without affecting fentanyl choice.

**Conclusions:** Results extend the range of conditions across which pain-related manipulations fail to increase opioid seeking in rats and suggest that enhanced opioid-addiction risk in humans with chronic pain involves factors other than enhanced opioid reinforcement and relapse.

## INTRODUCTION

Humans often take opioid drugs to reduce pain (Yaksh and Wallace 2018). Although the rates of iatrogenic (medically caused) addiction are low (~3%), this percentage still represents many cases (Epstein et al. 2018), and chronic pain-related factors increase the risk for opioid addiction (Epstein et al. 2018; Nazarian et al. 2021). Many patients with opioid addiction report that they experience chronic pain (Juurlink and Dhalla 2012; Kaye et al. 2017a; Kaye et al. 2017b), with ~60% of opioid users experiencing chronic pain (compared to 20% in general population) (Cicero et al. 2008; Dahlhamer et al. 2018). Considering the opioid overdose crisis, there is an urgent need to develop animal models to study behavioral and brain mechanisms underlying pain-related increases in risk for opioid addiction and to develop treatment strategies to decrease opioid use in users with comorbid pain.

One hypothesis for pain-related risk of opioid addiction is that pain states increase net opioid reinforcement by combining the positive reinforcing effects of opioid drugs with negative reinforcement of opioid-induced pain relief. However, the existing preclinical literature provides limited evidence for this hypothesis (Nazarian et al. 2021). Early studies reported that pain-related manipulations increase oral fentanyl consumption in a two-bottle choice procedure in rats (Colpaert et al. 1982; Colpaert et al. 2001). In contrast, a later study that used several pain-related manipulations found decreased acquisition of oral fentanyl consumption in mice (Wade et al. 2013). Additionally, inflammatory and neuropathic pain-related manipulations typically decrease the potency and/or effectiveness of opioids to maintain intravenous (i.v.) drug self-administration in rats (Hipolito et al. 2015; Lyness et al. 1989; Martin and Ewan 2008; Martin et al. 2007). Together, these studies do not support the notion that pain states increase opioid reinforcement (Nazarian et al. 2021).

However, one issue to consider in the studies cited above is that they may not have afforded sufficient opportunity for the subjects to learn that intravenous opioid self-administration can also relieve pain, in addition to positive reinforcement, and thus associate opioid self-administration with pain relief (negative reinforcement). In this regard, previous studies reported that non-contingent administration of mu opioid receptor agonists decreases the negative-reinforcing effects of noxious stimuli in avoidance tasks and the discriminative stimulus effects of noxious stimuli in discrimination tasks (Boada et al. 2016; Dykstra and McMillan 1977; Grilly et al. 1980; Harte et al. 2016; Thomas et al. 1992; Vierck et al. 2002). Based on these studies, we hypothesized that pain states can serve as negative reinforcer stimuli for contingent opioid self-administration. Therefore, in Experiments 1-2, we used different variations of the reinstatement model (Crombag et al. 2008; Shaham et al. 2003) and examined whether pairing an acute noxious stimulus with daily opioid self-administration would allow the rats to learn that opioid self-administration can relieve pain in a specific context. After extinction of the opioid-reinforced responding under pain-free conditions, we tested whether acute re-exposure to the noxious stimulus in the opioid self-administration context would reinstate opioid seeking or potentiate opioid priming-induced reinstatement. We chose intraplantar capsaicin and i.p. lactic acid as the acute noxious stimuli because they produce transient acute pain states (Gilchrist et al. 1996; Hohmann et al. 2005; Pereira Do Carmo et al. 2009) and thus would allow us to repeatedly pair an acute noxious stimulus with opioid self-administration.

A second issue to consider with the cited studies described above is that they primarily relied on single-operant self-administration procedures that generate rate-based measures of opioid self-administration. Rates of opioid self-administration can be reduced not only by selective decreases in reinforcing effects of the self-administered opioid, but also by nonselective motor or motivational impairments (Mello and Negus 1996; Negus and Banks 2011). Consistent with this latter possibility, many studies have reported that pain-related manipulations decrease self-administration of food (Cone et al. 2018; Martin et al. 2004), and electrical brain stimulation (Brust et al. 2016; Negus 2013). Thus, it is possible that pain-related decreases in operant responding would oppose and prevent increases in opioid self-administration due to pain-related increase in opioid reinforcement.

To address this concern, in Experiment 3, we examined the effect of pain-related manipulations on fentanyl vs. food choice. Choice procedures dissociate behavioral allocation (i.e. drug vs. food choice) and overall behavioral rate (i.e. total reinforcement rates maintained by both drug and food), and these two dependent variables allow for more refined interpretation of effects produced by experimental manipulations (Banks and Negus 2012; 2017). Specifically, selective changes in drug reinforcement are indicated by shifts in the drug-choice dose-effect curve, whereas nonselective motor or motivational impairment effects on operant responding are indicated by decreases in reinforcement rates. We predicted that if pain states increase opioid reinforcement, they would shift the opioid-choice dose-effect curve leftward, independent of a change in reinforcement rates. We chose i.p. lactic acid and intraplantar Complete Freund’s Adjuvant (CFA) as the noxious stimuli because this approach would allow us to compare the effects of intermittent vs. sustained models of inflammatory pain (Leitl et al. 2014) on fentanyl vs. food choice.

## MATERIALS AND METHODS

In the Supplemental Online Section, we describe the subjects (male and female rats), apparati, drugs, surgical procedures, pain-related manipulations, and the food and drug self-administration, reinstatement, and choice procedures used in Experiments 1-3. We also describe in this section the statistical analysis procedures. Below, we provide an overview and rationale of Experiments 1-3. We performed all experiments in accordance with the NIH Guide for the Care and Use of Laboratory Animals (8th edition), under protocols approved by the NIDA IRP Animal Care and Use Committee or the Virginia Commonwealth University Institutional Animal Care and Use Committee.

### Experiment 1: Effect of intraplantar capsaicin on reinstatement of fentanyl seeking

The goal of Experiment 1 was to determine whether a pain state (induced by exposure to intraplantar capsaicin) previously paired with fentanyl self-administration in a specific context would reinstate drug seeking after pain-free extinction of fentanyl-reinforced responding. We hypothesized that repeated pairing of the pain state with fentanyl self-administration would increase drug self-administration and later on reinstate opioid seeking after pain-free extinction, because the rats had learned that operant responding during training relieves pain (a negative reinforcement mechanism). We chose intraplantar capsaicin to activate cutaneous nociceptors, because this manipulation produces transient and repeatable nocifensive-related behaviors (Gilchrist et al. 1996; Hohmann et al. 2005). We used a variation of the ABA context-induced reinstatement procedure (Crombag et al. 2008; Crombag and Shaham 2002) and paired fentanyl self-administration with capsaicin exposure (plus brief isoflurane exposure to perform the intraplantar injection) in context A (Capsaicin context) and no capsaicin in context B (No-capsaicin context), in which the rats were not exposed to isoflurane and did not receive intraplantar vehicle injections (to minimize mild anesthesia exposure). We then extinguished responding in both contexts under pain-free conditions and tested whether capsaicin exposure in the previously capsaicin-paired context A would reinstate fentanyl seeking. We first determined the effect of intraplantar capsaicin on food self-administration (Experiment 1A) to identify capsaicin doses that do not cause non-selective deficits in operant responding. We subsequently used these doses in the fentanyl self-administration-reinstatement study (Experiment 1B).

In Experiment 1B (and 2B), we first trained rats for food self-administration for 4 sessions so that any effects of pain-related manipulations during opioid self-administration could not be attributed to general pain-induced deficits in operant learning. In all experiments, we used opioids at intravenous doses that are antinociceptive (Altarifi et al. 2015; Schwienteck et al. 2019a; Schwienteck et al. 2019b; Townsend et al. 2020; Townsend et al. 2019b). [Note: We used male rats in Experiment 1 and both male and female rats in Experiments 2-3 but did not power the latter experiments to detect sex differences. Our intention was to perform follow-up experiments to detect sex differences if the effect of the pain-related manipulations on reinstatement and choice were positive.]

### Experiment 2: Effect of i.p. lactic acid on reinstatement of heroin seeking

In Experiment 1 we found that capsaicin had no effect on fentanyl self-administration and did not reinstate fentanyl seeking. However, a limitation of this experiment was that capsaicin exposure required mild anesthesia before the self-administration and reinstatement sessions, which may interfere with the rat’s behavior. Another limitation in Experiment 1 was the potential for tolerance to the nociceptive effects of capsaicin after repeated exposure (Lundberg and Saria 1983), resulting in decreased efficacy of capsaicin to induce pain-related states during self-administration training and reinstatement testing. Therefore, in Experiment 2, we used a different acute pain-related manipulation, daily i.p. injections of lactic acid, which does not require anesthesia (Negus 2013; Pereira Do Carmo et al. 2009) and for which tolerance does not develop after repeated exposure (Legakis et al. 2020; Miller et al. 2015). Furthermore, because the context manipulation in Experiment 1 was ineffective, we simplified the experimental design and eliminated the context in which a noxious stimulus was absent. We also switched from fentanyl to heroin, which allowed us to compare reinstatement induced by a noxious stimulus to reinstatement induced by drug priming and to determine whether the noxious stimulus would increase drug priming-induced reinstatement. [Note: In several pilot studies, we were unable to observe reliable fentanyl priming-induced reinstatement across multiple doses and routes of administration.] As in Experiment 1, we first determined the effect of the pain-related manipulation on food self-administration (Experiment 2A) to verify that at the dose range used in the heroin self-administration-reinstatement study (Experiment 2B), lactic acid does not cause non-selective deficits in operant responding.

In Experiment 2B, we first trained rats on food self-administration for 4 sessions so that any effects of pain-related manipulations during opioid self-administration could not be attributed to general pain-induced deficits in operant learning.

### Experiment 3: Effect of i.p. lactic acid and intraplantar CFA on choice fentanyl vs. food choice

In Experiments 1-2 we found no evidence that acute pain-related manipulations reinstate opioid seeking. A limitation of these experiments is the potential confounding factor of non-selective pain-related suppression of operant responding (Mello and Negus 1996; Negus and Banks 2011). To address this concern, we examined the effect of pain-related manipulations on choice between fentanyl and food, which allowed us to independently measure behavioral allocation (i.e., opioid choice) and behavioral rate (i.e., rate of reinforcement). We tested the effect of repeated i.p. lactic acid injections (Experiment 3A) and intraplantar CFA (Experiments 3B-C) on fentanyl vs. food choice.

## RESULTS

The goal of Experiment 1A and 2A (Fig. S1A and S2A) was to determine the effect of intraplantar injections of capsaicin or i.p injections of lactic acid on food self-administration. Based on these experiments, we used the optimal doses for Experiments 1B and 2B where we determined the effect of these pain-related manipulations on reinstatement of opioid (fentanyl or heroin) seeking after pairing capsaicin or lactic acid exposure with opioid self-administration. The full statistical analyses are provided in Supplemental Table S1.

### Capsaicin decreased food self-administration

The rats acquired food self-administration (data not shown). Repeated capsaicin (50 μg) injections had no effect on food self-administration (Fig. S1B, p values>0.05). However, in a subsequent dose-response determination capsaicin significantly decreased active lever presses (Fig. S1C). The analysis, which included the within-subjects factors of Capsaicin dose (0, 50, or 100 μg) and Lever (inactive, active) showed significant effect of Capsaicin dose (F_1,10_=9.5, p=0.005) and interaction between the two factors (F_1,10_=10.3, p=0.004). Capsaicin also decreased the number of food rewards earned, but this effect did not reach statistical significance (main effect of Capsaicin dose: (F_1,10_=3.5, p=0.07).

### Capsaicin had no effect on fentanyl self-administration or reinstatement

The goal of Experiment 1B (timeline Fig. 1A) was to determine whether exposure to capsaicin in a previously pain-associated context would reinstate fentanyl seeking after extinction.

**Figure 1.**
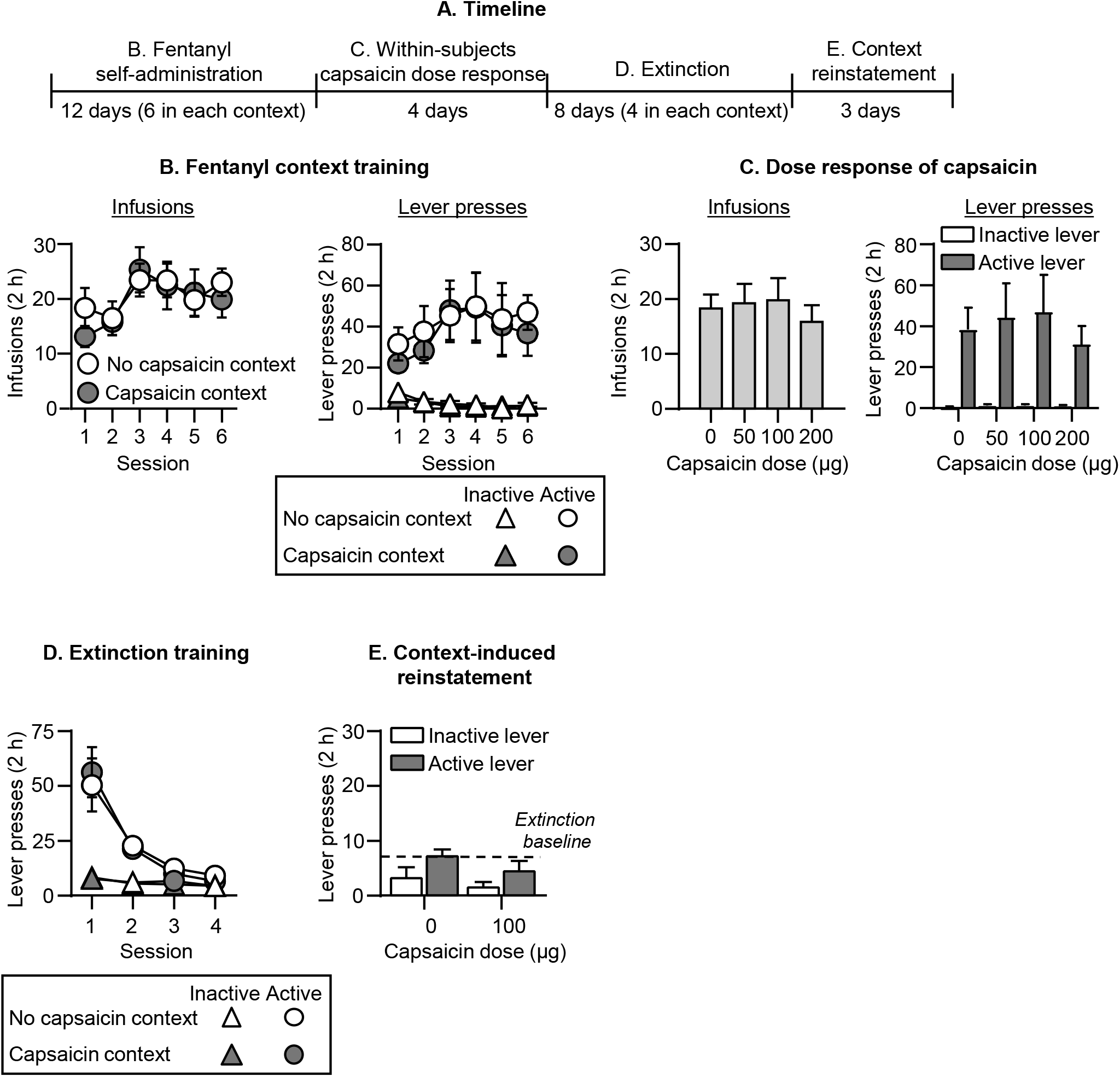
Effect of intraplantar capsaicin context on fentanyl seeking. **(A)** Experimental timeline of Experiment 1B. **(B)** <u>Fentanyl self-administration:</u> No capsaicin and Capsaicin contexts: Number of fentanyl infusions (2.5 μg/kg/infusion) during the 2-h sessions after exposure in the No capsaicin context or 100 μg capsaicin (intraplantar hindpaw injection under light isoflurane anesthesia) in the Capsaicin context (n=11, within-subjects design). **(C)** <u>Fentanyl self-administration: Capsaicin dose-response:</u> Number of fentanyl infusions during the 2-h sessions after 0 (vehicle), 50, 100, and 200 μg capsaicin injections (n=11, within-subjects design). **(D)** <u>Extinction in No capsaicin and Capsaicin contexts:</u> Number of inactive and active lever presses during the 2-h extinction sessions in the No capsaicin context and in the Capsaicin context (n=11, within-subjects design). **(E)** <u>Reinstatement in Capsaicin context:</u> Number of inactive and active lever presses during the 2-h sessions after 0 (vehicle; n=6) or 100 μg capsaicin injection (n=5) in the Capsaicin context. Number of active lever presses after in the No capsaicin context (n=11) is depicted as a baseline dotted line. Data are mean□±SEM.

#### Self-administration training

Repeated capsaicin (100 μg) injections had no effect on fentanyl self-administration (Fig. 1B). The analysis of fentanyl infusions, which included the within-subjects factors of Context (No Capsaicin, Capsaicin) and Session (1-6), showed a significant effect of Session (F_5,50_=5.6, p<0.001) but not Context or interaction. Capsaicin also had no effect on fentanyl infusions or lever presses during a subsequent dose-response (0, 50, 100, or 200 μg) determination (Fig. 1C, Fig. S4A for individual data).

#### Extinction training

The rats extinguished responding for fentanyl over sessions in both the previously Capsaicin-paired context and the No-Capsaicin context, and the context had no effect on extinction (Fig. 1D). The analysis of lever pressing, which included the within-subjects factors of Session, Context (No Capsaicin, Capsaicin), and Lever showed significant effects of Session x Lever (F_3,30_=15.5, p<0.001) but no effect of Context or interactions between Context and the other factors.

#### Reinstatement

Capsaicin injections in the previously capsaicin-paired training context had no effect on reinstatement of fentanyl seeking (Fig. 1E, Fig. S4B for individual data). The analysis of lever presses, which included the between-subjects factor of Capsaicin dose (0, 100 μg) and Lever, showed a main effect of Lever (F_1,9_=9.4, p=0.013) but not Capsaicin dose or interaction.

### Lactic acid decreased food self-administration

Both concentrations of lactic acid (0.9% and 1.8%) decreased food self-administration (Fig. S2B). The analysis of number of rewards, which included the within-subjects factor of Lactic acid concentration (0, 0.9%, or 1.8%), showed a significant effect (1.8%: F_1,7_=89.6, p<0.001; 0.9%: F_1,7_=6.6, p=0.037). Lactic acid (0, 0.9, 1.35, 1.8%) also decreased food self-administration under the progressive ratio schedule (Fig. S2C). Lactic acid increased Pain-related behaviors and decreased No-Pain-related behaviors (Fig. S2D). The analysis of number of behaviors (see Supplemental Methods), which included the within-subjects factors of Behavior type (No pain, Pain) and Lactic acid concentration, showed a significant interaction between the two factors (F_3,21_=36.2, p<0.001).

### Lactic acid had no effect on heroin self-administration or reinstatement

The goal of Experiment 2B (timeline Fig. 2B) was to determine whether re-exposure to lactic acid (previously paired with heroin self-administration) would reinstate heroin seeking after extinction. We injected either sterile water vehicle or 0.9% lactic acid daily during the heroin self-administration phase.

**Figure 2.**
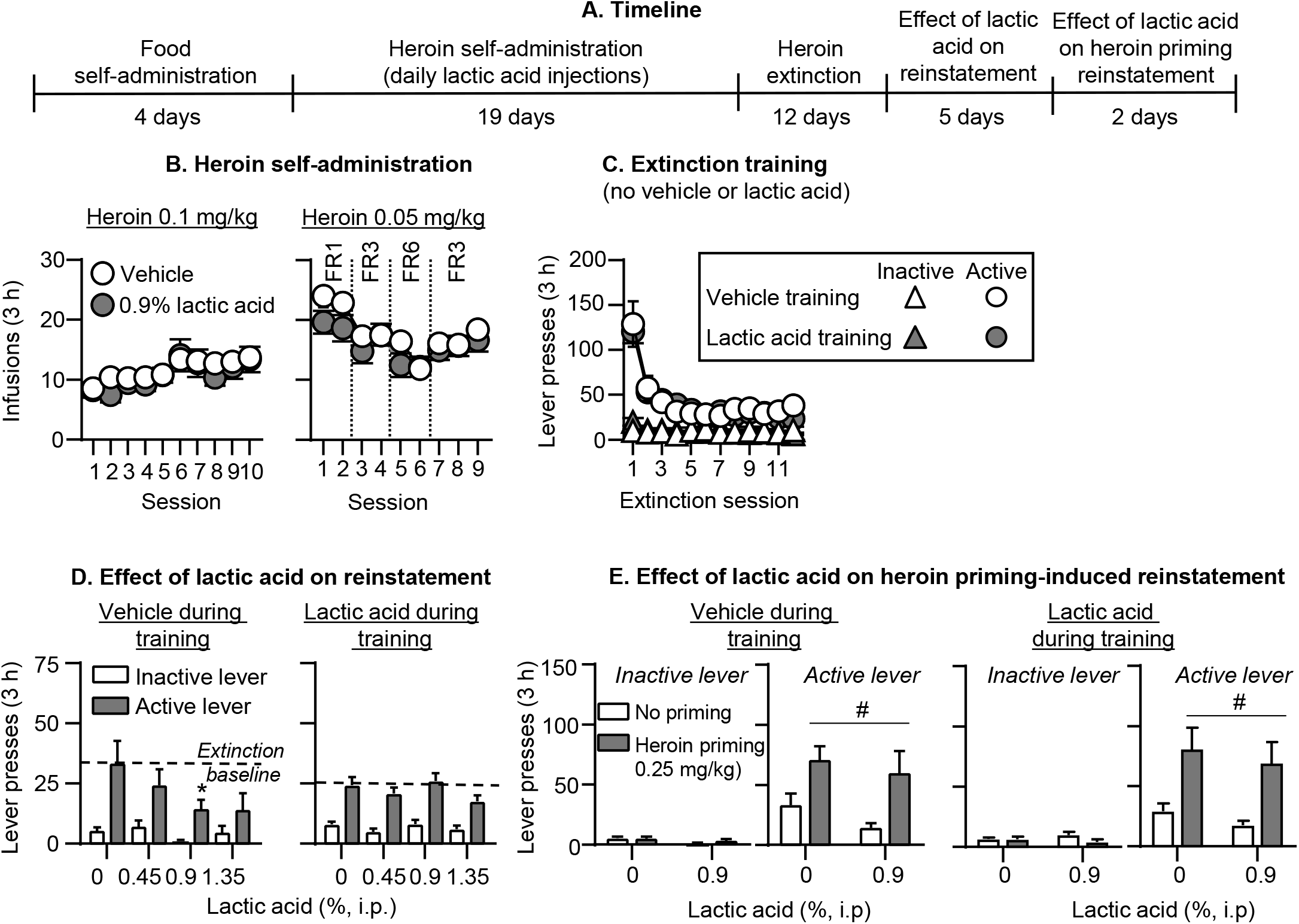
Effect of i.p. lactic acid on reinstatement of heroin seeking. **(A)** Experimental timeline of Experiment 2B. **(B)** <u>Heroin self-administration:</u> Number of heroin infusions (0.1 and 0.05 mg/kg/infusion) during the 3-h self-administration sessions (vehicle group, n=7; lactic acid group, n=13). **(C)** <u>Extinction:</u> Number of inactive and active lever presses during the 3-h extinction sessions (vehicle group, n=6; lactic acid group, n=13). **(D)** <u>Effect of lactic acid on reinstatement:</u> Number of inactive and active lever presses during the 3-h sessions after lactic acid injections (0, 0.45, 0.9, 1.35%, i.p.). Number of active lever presses over the last 3 extinction sessions is depicted as a baseline dotted line (vehicle group during training: n=6; lactic acid group during training, n=13) **(E)** <u>Effect of lactic acid on heroin priming-induced reinstatement:</u> Number of inactive and active lever presses during the 3-h sessions (left: vehicle training condition, right: lactic acid training condition). Rats received either no heroin priming (data re-graphed from 2D) or 0.25 mg/kg heroin injections (s.c.) and either 0% or 0.9% lactic acid (i.p.). (vehicle group during training: n=6 saline priming, n=6 heroin priming; within-subjects design; lactic acid group during training: n=6 saline priming, n=7 heroin priming; between-subjects design). * Different from 0% lactic acid, p<0.05. # Different from no heroin priming, p<0.05. Data are mean□±SEM.

#### Self-administration and extinction

Repeated lactic acid injections had no effect on heroin self-administration under the different fixed ratio schedule requirements (Fig. 2B). The analysis of heroin infusions, which includes the between-subjects factor of Training condition (Vehicle, 0.9% lactic acid) and the within-subjects factor of Session showed a significant effect of Session (F_18,324_=12.9, p<0.001) but not Training condition or interaction. We were initially concerned that lactic acid was interfering with acquisition of heroin self-administration and gave all rats an additional heroin training session without lactic acid injections after session 4, and 2 rats an additional heroin training session without lactic acid injections after session 7. Responding from these sessions was not significantly different than adjacent sessions (p values>0.05) and we do not present or analyze data from these extra sessions. [Note: our original intention was to maintain the rats on an FR6 schedule toward the end of training, and during the extinction and reinstatement phases, but several rats did not maintain their heroin intake under this schedule, thus we switched all rats back to the FR3 schedule.] Capsaicin also had no effect on within-session heroin self-administration (see representative results from Session 12 (FR1) and Session 19 (FR3) in Fig. S3A and statistical analysis results in Table S1).

During the extinction phase, the rats decreased responding on the previously active lever presses (Fig. 2C). The analysis of number of lever presses, which included the between-subjects factor of Training condition and the within-subjects factors of Session and Lever, showed an interaction of Session x Lever (F_11,187_=25.7, p<0.001) but no significant effect of Training condition or interaction with this factor.

#### Reinstatement

Lactic acid injections did not reinstate heroin seeking after extinction but decreased active lever presses in the vehicle but not lactic acid training group (Fig. 2D, Fig. S5A for individual data). The analysis of lever presses, which included the between-subjects factor of Training condition (vehicle, lactic acid) and the within-subjects factors of Lactic acid concentration and Lever, showed significant interactions of Training condition x Lactic acid concentration (F_3,51_=4.5, p=0.007) and Training condition x Lactic acid concentration x Lever (F_3,51_=2.9, p=0.04). Post-hoc analysis within each training condition showed a significant interaction of Lactic acid concentration x Lever for the vehicle (F_3,15_=3.7, p=0.04) but not lactic acid training condition.

In subsequent tests for heroin priming-induced reinstatement, lactic acid (0.9%) injections had no significant effects (Fig. 2E, Fig. S5B for individual data). To represent baseline no heroin priming responding with or without 0.9% lactic acid, we used data from Fig. 2D in Fig. 2E. In rats that received vehicle injections during training, the analysis of lever presses, which included the within-subjects factors of Lactic acid concentration (0 or 0.9%), Heroin priming (no priming, 0.25 mg/kg), and Lever, showed a significant effect of Heroin priming (F_1,5_=8.9, p=0.03) but not Lactic acid concentration or interaction. In rats that received 0.9% lactic acid injections during the training phase, the analysis of lever presses, which included the between-subjects factor of Lactic acid concentration and the within-subjects factors of Heroin priming and Lever, showed a significant effect of Heroin priming (F_1,11_=12.0, p=0.005) but not Lactic acid concentration or interaction.

### Lactic acid had no effect on fentanyl vs. food choice

The goal of Experiment 3 (timeline Fig. 3A) was to test the effect of noxious stimuli on fentanyl vs. food choice, which allowed us to distinguish between effects on opioid reinforcement and behavioral suppression. Following stable fentanyl vs. food choice behavior, we determined the effect of lactic acid injections on fentanyl vs. food choice. We injected lactic acid repeatedly to provide an opportunity for the rats to shift their choice behavior in response to learning without changing the contingencies associated with pain produced by the lactic acid and pain relief produced by self-administered fentanyl.

**Figure 3.**
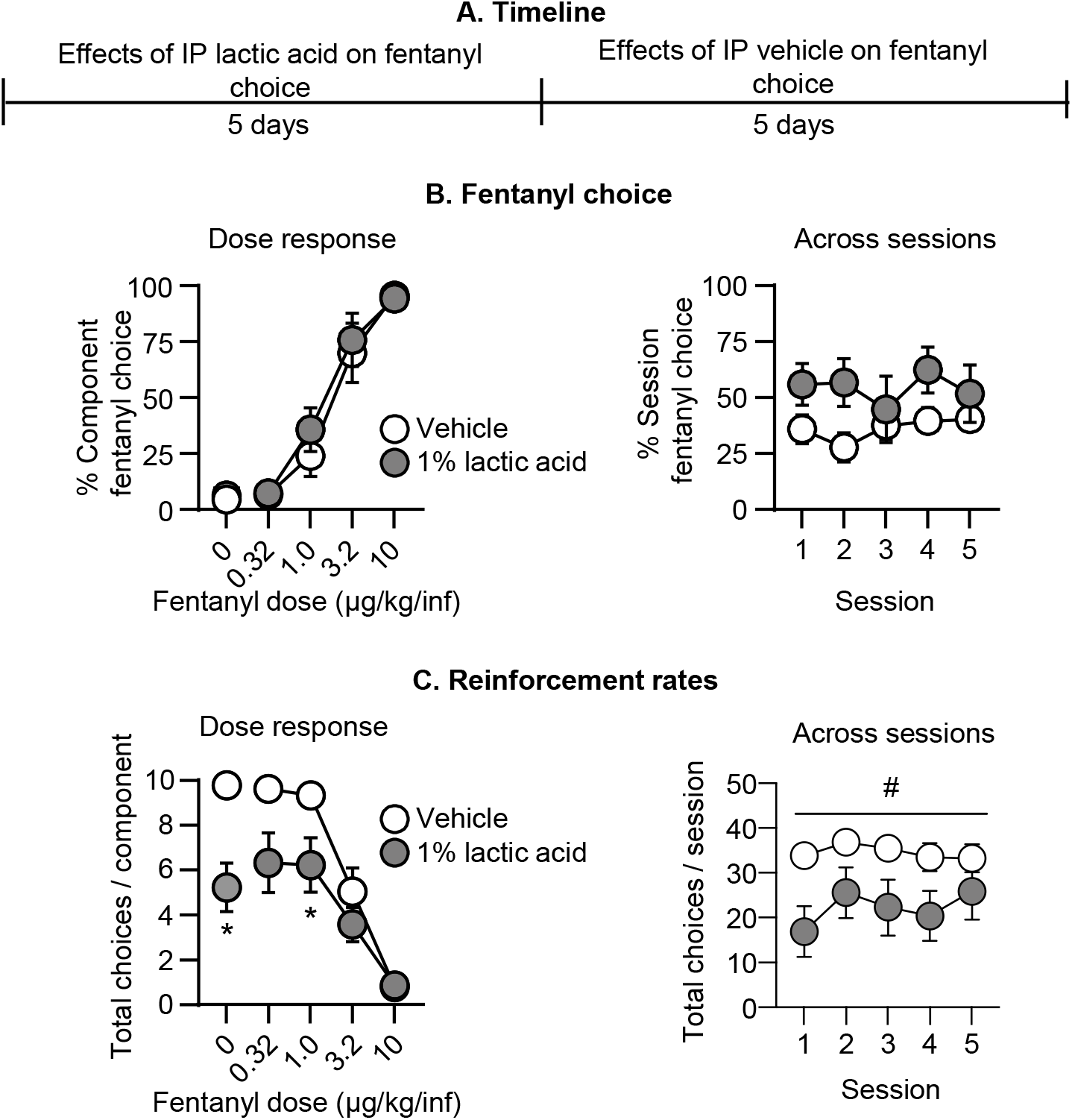
Effect of repeated i.p. lactic acid on fentanyl vs. food choice. **(A)** Experimental timeline of Experiment 3A. **(B)** <u>Fentanyl choice:</u> Percent fentanyl choice across within-session fentanyl unit doses in rats injected i.p. with vehicle or 1% lactic acid (n=8, counterbalanced across rats). <u>Left:</u> Averaged across the five days of treatment and shown as a function of fentanyl unit dose. Right: Averaged across fentanyl unit dose and shown as a function of session for the five days of treatment. **(C)** <u>Reinforcement rates:</u> Total choices completed per component (maximum of 10 choices per component) across the within-session reinforcement-rate dose-response function. <u>Left:</u> averaged across the five days of treatment and shown as a function of fentanyl unit dose. <u>Right:</u> Averaged across fentanyl unit dose and shown as a function of session for the five days of treatment. * Difference from vehicle treatment at a given fentanyl unit dose, p<0.05. # Significant main effect of lactic acid, p<0.05. Data are mean□±SEM.

#### Fentanyl choice

When we collapsed data across treatment days, repeated pretreatment with 1% lactic acid did not significantly affect the fentanyl choice dose-effect curve relative to vehicle pretreatment (Fig. 3B). The analysis of percent fentanyl choice, which included the within-subjects factors of Fentanyl unit dose (0-10 μg/kg/infusion) and Lactic acid (0, 1%), showed a significant effect of Fentanyl unit dose (F_1.6, 11.1_=50.7, p<0.0001) but not Lactic acid or interaction (p values>0.05). Similarly, when we collapsed data across fentanyl unit dose, lactic acid did not affect total session fentanyl choice across the five consecutive days of pretreatment (Fig. 3B). The analysis showed no effect of Session, Lactic acid, or interaction (see Table S1).

#### Reinforcement rates

Repeated 1% lactic acid injections significantly decreased the total number of choices completed per component (Fig. 3C). The analysis showed a significant interaction of Fentanyl unit dose and Lactic acid (F_1.8, 12.5_=9.0, p=0.0045), with significant decreases during availability of 0 and 1.0 μg/kg/inf fentanyl, when rats primarily chose food. When data were collapsed across fentanyl unit doses (Fig. 3C), the analysis showed a significant effect of Lactic acid (F_1.0, 7.0_=17.2, p=0.004) but no effect of Session or interaction. Thus, repeated lactic acid failed to alter fentanyl choice despite producing a repeatable decrease in reinforcement rates.

### CFA had no effect on fentanyl vs. food choice

In the same rats used in Experiment 3A, we next tested the effect of intraplantar CFA as a model of sustained inflammatory pain on fentanyl vs. food choice (timeline Fig. 4A). We also determined the effect of CFA on mechanical sensitivity and paw width as additional measures of pain-related behavior and inflammation.

**Figure 4.**
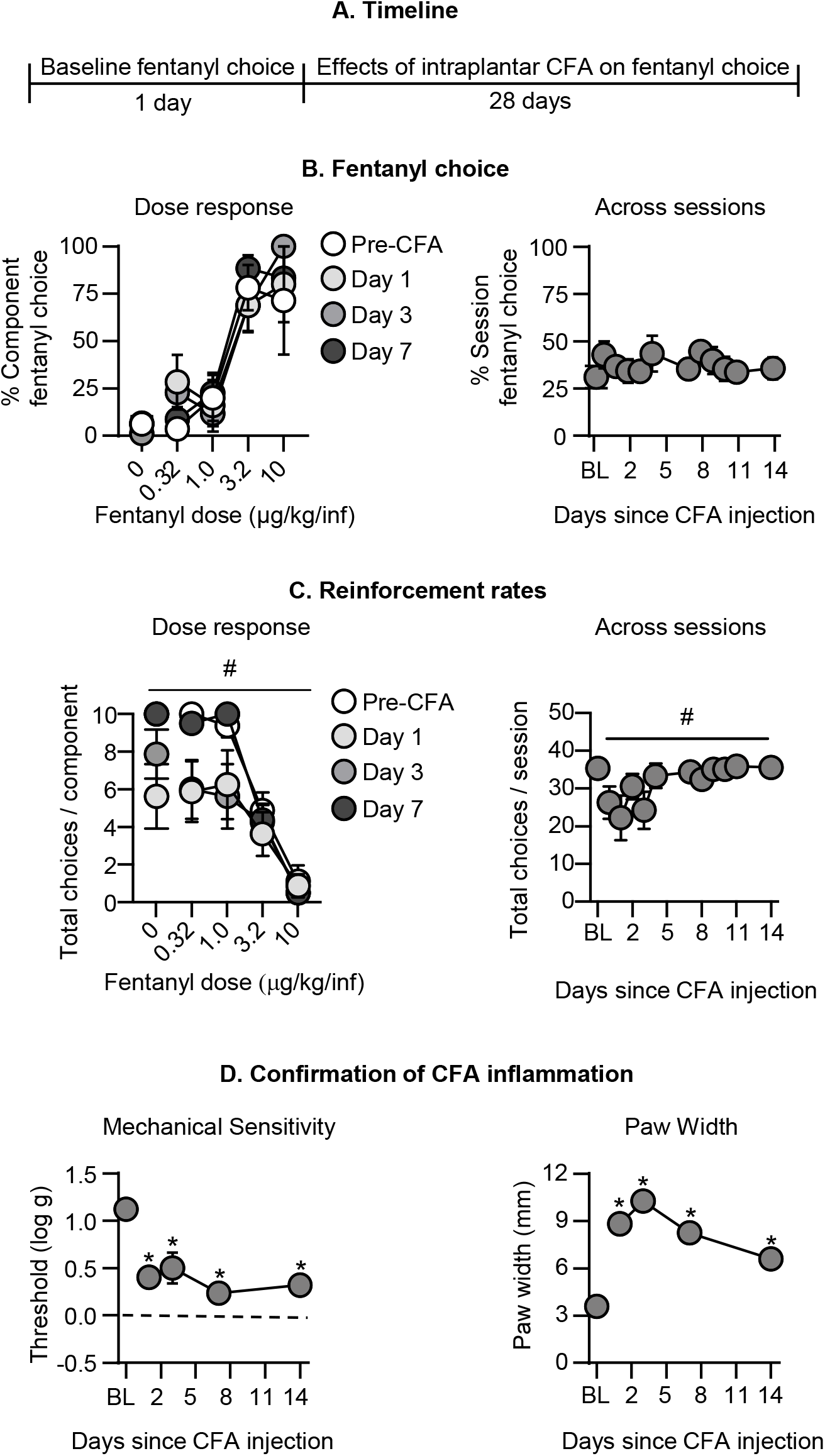
Effect of intraplantar CFA on fentanyl vs. food choice. **(A)** Experimental timeline of Experiment 3B. **(B)** <u>Fentanyl choice:</u> Percent fentanyl choice across within-session fentanyl unit doses in rats with an intraplantar injection of CFA (n=8). <u>Left:</u> Within-session fentanyl-vs.-food choice dose-response function on Days 1, 3, and 7 following intraplantar CFA administration. <u>Right:</u> Session percent fentanyl choice across the first 14 days following intraplantar CFA, averaged across fentanyl unit dose. **(C)** <u>Reinforcement rates:</u> Total choices completed per component (maximum of 10 choices per component) across the within-session reinforcement-rate dose-response function. <u>Left:</u> Within-session fentanyl-vs.-food choice dose-response function on Days 1, 3, and 7 following intraplantar CFA administration. <u>Right:</u> Total choices per session across the first 14 days following intraplantar CFA, averaged across fentanyl unit dose. **(D)** <u>Confirmation of CFA inflammation: Left:</u> Effect of intraplantar CFA on mechanical sensitivity (Threshold (log g) for paw withdrawal from von Frey filaments). <u>Right:</u> Effect of intraplantar CFA on paw width (mm). * Difference from baseline at a given timepoint, p<0.05. # Significant main effect of time since intraplantar CFA, p<0.05. Data are mean□±SEM.

#### Fentanyl choice

CFA had no effect on percent fentanyl choice across selected timepoints during the first week after its administration (Fig. 4B). The analysis, which included the within-subjects factors of Fentanyl unit dose and Session (Baseline, 1-14) showed a significant effect of Fentanyl unit dose (F_1.6, 11.0_=36.8, p<0.0001) but not Session or interaction. When collapsed across fentanyl unit doses, CFA had no effect on fentanyl choice through 14 days post-CFA administration (Fig. 4B). We also did not detect an effect of CFA on fentanyl choice 21 or 28 days after the CFA injection (data not shown).

#### Reinforcement rates

CFA injection significantly decreased the number of choices completed per component (Fig. 4C). The analysis, which included the within-subjects factors of Fentanyl unit dose and Session, showed significant effects of Fentanyl unit dose (F_1.9, 13.0_=73.9, p<0.0001) and Session (F_1.3, 8.9_=6.0, p=0.03) but no interaction. When data were collapsed across fentanyl unit doses, CFA decreased choices per session (Fig. 4C; F_2.3, 16.0_=4.5, p=0.02).

#### Confirmation of CFA inflammation

These significant changes in rates of operant responding corresponded with sustained increases in mechanical sensitivity (Fig. 4D; F_2.1, 14.5_=17.5, p=0.0001) and paw width (Fig. 4D; F_2.4, 16.7_=100.9, p<0.0001) for days 1-14 after CFA. We also measured significant mechanical hypersensitivity and paw swelling that persisted until Day 28 after CFA (data not shown). Thus, CFA failed to alter fentanyl choice despite producing transient decreases in reinforcement rates and sustained mechanical hypersensitivity and paw swelling.

### Effect of manipulating fixed ratio requirements on fentanyl vs. food choice

Because lactic acid and CFA produced little change in fentanyl choice despite evidence for significantly decreasing reinforcement rates, we performed a follow-up experiment (timeline Fig. 5A) to examine sensitivity of choice to a change in response requirements as a positive control.

**Figure 5.**
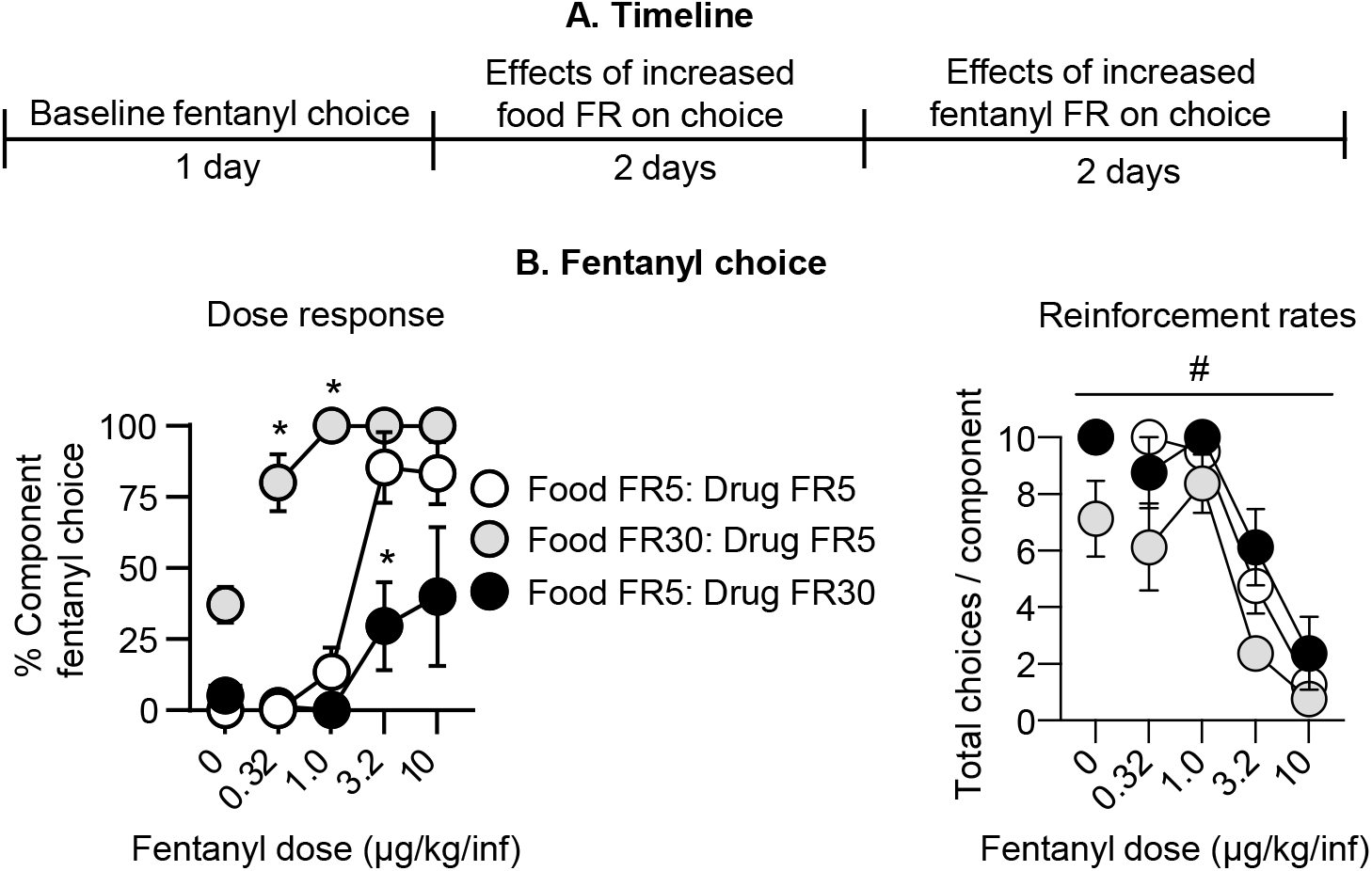
Effect of FR manipulations on fentanyl vs. food choice. **(A)** Experimental timeline of Experiment 3C. **(B)** <u>Fentanyl choice:</u> Percent fentanyl choice across within-session fentanyl unit doses following FR response requirement manipulations in rats after completion of Experiment 3B. <u>Left:</u> Within-session fentanyl-vs.-food choice dose-response function. <u>Right:</u> Total choices completed per component (maximum of 10 choices per component) across the within-session dose-response function. * Difference from baseline schedule (i.e., Food FR5: Drug FR5) at a given unit fentanyl dose, p<0.05. # Significant main effect of FR manipulation, p<0.05. Data are mean□±SEM.

#### Fentanyl choice

Independently increasing the response requirement for food or fentanyl significantly affected fentanyl choice. Specifically, selectively increasing the food FR requirement while holding the fentanyl FR requirement at the original value (food FR 30: fentanyl FR 5) reduced food choice and reciprocally increased fentanyl choice. Conversely, holding the food FR requirement at the original value while selectively increasing the fentanyl FR requirement (food FR 5: fentanyl FR 30) increased food choice and reduced fentanyl choice (Fig. 5B). The analysis of fentanyl choice, which included the within-subjects factors of Fentanyl unit dose and FR requirement showed a significant interaction of Fentanyl unit dose x FR requirement (F_3.7, 22.1_=7.9, p=0.0005).

#### Reinforcement rates

In line with this analysis, we also detected a significant main effect of the FR requirement manipulation (F_1.6, 11.0_=12.4, p=0.002) and fentanyl unit dose (F_2.1, 14.9_=85.0, *p*<0.0001) on the number of choices completed per component but no interaction (Fig. 5B). Thus, while fentanyl choice was insensitive to lactic acid or CFA as intermittent or sustained pain-related manipulations, it could be significantly increased or decreased by increasing the FR requirement for food or fentanyl.

## DISCUSSION

We evaluated the effect of pain-related manipulations on opioid self-administration, reinstatement of opioid seeking, and opioid choice. We found that the inflammatory pain models we used (intraplantar capsaicin, i.p. lactic acid, inraplantar CFA) decreased operant responding for food and produced pain-related behaviors and measures of inflammation. However, despite this evidence for pain states, neither capsaicin nor lactic acid altered ongoing opioid self-administration or promoted reinstatement of opioid seeking. Additionally, neither lactic acid nor CFA increased opioid vs. food choice. Our results extend the range of conditions across which pain-related manipulations fail to increase metrics of opioid reinforcement and seeking in rats. To the degree the rat relapse and choice models we used here translate to humans (Banks and Negus 2017; Venniro et al. 2020), our results do not support the hypothesis that increased risk for opioid addiction in patients with pain is due to pain-related increases in opioid reinforcement or opioid seeking.

### Pain-related effects of capsaicin, lactic acid, and CFA

The pain-related behavioral effects of capsaicin, lactic acid, and CFA observed here agree with results from previous studies (Negus 2019). Thus, capsaicin’s effectiveness to decrease operant responding for food in our study is consistent with its effectiveness to produce nocifensive behaviors like paw-flinching (Ji et al. 2006), hypersensitive withdrawal responses from thermal and mechanical stimuli (Gilchrist et al. 1996; Hohmann et al. 2005), and hypersensitivity to thermal punishment of food self-administration (Neubert et al. 2006). Similarly, lactic acid’s effectiveness to increase pain-related behaviors observed here is consistent with previous evidence that lactic acid produces stretching/writhing behaviors (Pereira Do Carmo et al. 2009), facial grimace (Langford et al. 2010), and decreased operant responding maintained by food (Cone et al. 2018), or electrical brain stimulation (Negus 2013). Lastly, we found that CFA produced sustained mechanical hypersensitivity and paw inflammation and transiently (~3-day) decreased food self-administration during early components of daily operant sessions when rats chose primarily food. These results agree with evidence that CFA produces sustained mechanical hypersensitivity and swelling (Leitl et al. 2014; Stein et al. 1988), as well as transient facial grimace (Sotocinal et al. 2011) and depression of operant responding maintained by electrical brain stimulation (Leitl et al. 2014) or food (Negus et al. 2020).

Together, these results indicate that the pain-related manipulations in our study were sufficient to produce pain states, allowing us to examine their impact on opioid self-administration, reinstatement, and choice. Furthermore, the dose range of intravenous fentanyl and heroin in our study has been shown to be antinociceptive in previous studies (Schwienteck et al. 2019b; Townsend et al. 2019a; Townsend et al. 2020). We also reported that s.c. fentanyl alleviated i.p. lactic acid-induced pain behaviors with ED50 values of 4.0-8.0 μg/kg (Altarifi et al. 2015). Given that fentanyl is ≥5x more potent after i.v. than s.c. administration (see (Schwienteck et al. 2019b; Townsend et al. 2019a)), these data provide further evidence that the i.v. fentanyl doses used here are likely to be antinociceptive and suggest the potential for these pain states to function as negative reinforcers for self-administration of these opioid doses.

### Lack of effect of pain-related manipulations on opioid self-administration

We found that neither capsaicin nor lactic acid altered fentanyl or heroin self-administration. Additionally, neither repeated lactic acid nor CFA altered rates of fentanyl choice when high doses of fentanyl were available and the rats primarily chose fentanyl. However, the data in Experiments 1-2 should be interpreted with caution, because we only determined the effect of the pain-related manipulations on a single fentanyl unit dose or two heroin unit doses. To address this issue, we included a full dose-effect analysis to allow for interpretation of fentanyl’s reinforcing effects in a rate-independent choice procedure in Experiment 3.

Our findings agree with previous studies showing that, once self-administration has been established for high opioid unit doses (doses on the descending limb of opioid dose-effect curve under low-rate fixed-ratio schedules), this self-administration is resistant to reductions by pain states that decrease operant responding of non-opioid reinforcers (see above) (Hipolito et al. 2015; Martin et al. 2007). This resilience of high-dose opioid self-administration in the presence of inflammatory and neuropathic pain models may be related to opioid analgesia and the negative reinforcing effects of pain because the self-administered opioid doses produce antinociception when administered non-contingently (Nazarian et al. 2021). However, there was no evidence in our study or the Martin et al. (2007) study that pain states increase opioid self-administration in a way that might suggest a net increase in opioid reinforcement. Furthermore, although Hipolito et al. (2015) reported that CFA increased high-dose heroin intake under an FR schedule, CFA had no effect on self-administration of the same heroin dose under a progressive-ratio schedule.

In contrast to the weak effects of pain-related manipulations on high-dose opioid self-administration, pain-related manipulations have been reported to decrease self-administration of lower unit doses of opioids (doses on the ascending limb or peak of the self-administration dose-effect curve under low-rate fixed ratio schedules) (Hipolito et al. 2015; Martin et al. 2007). Moreover, pain states have been reported to decrease acquisition of opioid self-administration when the pain state is implemented before initiation of self-administration training (Lyness et al. 1989; Wade et al. 2015), but see (Colpaert et al. 2001). In our study, repeated lactic acid did not alter acquisition of heroin self-administration (see Fig. 2B). Furthermore, the negative findings in our study contrast with the effectiveness of experimenter-delivered non-contingent opioids to decrease the negative reinforcing efficacy of noxious stimuli in avoidance tasks and discriminative stimulus effects of noxious stimuli in discrimination tasks (Boada et al. 2016; Dykstra and McMillan 1977; Grilly et al. 1980; Harte et al. 2016; Thomas et al. 1992; Vierck et al. 2002).

### Lack of pain-related manipulation effects on reinstatement of opioid seeking

The published studies described above provide little evidence that pain states increase opioid reinforcement. However, to our knowledge, the effect of pain-related manipulations on reinstatement after prior exposure to pain-related stimuli during ongoing opioid self-administration has not been determined. We hypothesized that, by explicitly pairing noxious stimuli with opioid self-administration, rats will learn that opioid intake decreases pain during self-administration, and subsequent exposure to the noxious stimuli will induce reinstatement of opioid seeking after pain-free extinction of opioid-reinforced responding. Our results do not support this hypothesis: neither capsaicin nor lactic acid exposure induced pain-related reinstatement of fentanyl or heroin seeking, respectively. Additionally, lactic acid decreased baseline extinction responding and had no effect on heroin priming-induced reinstatement. However, unlike the results for reinstatement induced by capsaicin and lactic acid where we used multiple concentrations, the negative results with heroin priming should be interpreted with caution because we only used a single concentration of lactic acid and a single heroin priming dose.

Regarding the negative results with capsaicin, two other alternative interpretations should be considered. The first limitation is that, unlike the lactic acid experiment where the rats received 19 sessions of pairing lactic acid exposure with heroin self-administration, the rats in the capsaicin experiment only received 6 sessions of pairing capsaicin exposure with fentanyl self-administration. This number of pairings may not be sufficient for the rats to learn that fentanyl self-administration can relieve pain. The second limitation is potential desensitization of C-fibers after repeated capsaicin injections (Lundberg and Saria 1983). However, while we cannot rule out this possibility, it is relatively unlikely because in Experiment 1A, capsaicin decreased food responding after repeated injections (Fig. S1 dose response). To address these limitations, in Experiment 2, we modified the experimental design and increased the number of pairings between pain exposure and opioid self-administration. We used lactic acid as a noxious stimulus because it can be administered repeatedly without developing tolerance to its pain-related decrease on operant responding (Legakis et al. 2020; Miller et al. 2015).

Finally, our negative results indicate that, unlike some acute stress manipulations like intermittent footshock and brain injections of the stress hormone corticotropin-releasing factor (Mantsch et al. 2016; Shaham et al. 2000), the acute pain-related manipulations used in our study are not effective stimuli for reinstatement of opioid seeking.

### Lack of pain-related manipulation effects on opioid vs. food choice

Previous studies on the effect of pain-related stimuli on intravenous opioid self-administration relied almost exclusively on single-operant self-administration procedures that generate rate-based measures of opioid reinforcement (Nazarian et al. 2021). However, in these types of procedures, pain-related depression of general behavior can obscure more selective pain-related changes in opioid reinforcement associated with opioid-induced pain relief. Thus, as a second strategy to expand our assessment of pain-related manipulation effects on opioid-taking behaviors, we examined the effects of repeated lactic acid and of CFA on fentanyl vs. food choice. Choice procedures allow for dissociation between selective treatment effects on opioid reinforcement (indicated by changes in % opioid choice) from nonselective treatment effects on motor competence and motivation (indicated by changes in reinforcement rate) (Banks and Negus 2012).

We found that neither repeated lactic acid nor CFA altered fentanyl choice, but both decreased reinforcement rates. These findings suggest that these pain-related manipulations produced nonselective behavioral disruption without altering opioid reinforcement. The lack of effect of lactic acid or CFA on fentanyl choice is not due to lack of sensitivity of our procedure, because changes in the fentanyl or food FR values did alter fentanyl choice. Additionally, we previously reported changes in fentanyl vs. food choice after other manipulations that include spontaneous opioid withdrawal and treatment with naltrexone maintenance or a fentanyl vaccine (Townsend et al. 2019a). Our finding that pain-related manipulations did not alter opioid choice is also consistent with the lack of effect of pain-related manipulations on the rewarding effects of opioids in studies using intracranial self-stimulation or place-conditioning procedures (Moerke and Negus 2019; Nazarian et al. 2021). Finally, our data are consistent with other evidence for effectiveness of lactic acid and CFA in rats to produce general depression of other behaviors, including feeding, wheel running, and burrowing (Andrews et al. 2012; Kandasamy and Morgan 2020; Kwilasz and Negus 2012; Stevenson et al. 2011), and decreases in self-administration of non-drug reinforcers (see above).

### Clinical implications and conclusions

The literature on opioid intake and relapse in patients with chronic pain is mixed (Nazarian et al. 2021). Some studies with patients with pain report increased opioid craving but no correlation between pain severity and positive urine toxicology results or treatment outcomes (Fox et al. 2012; Kantor et al. 1980; Potter et al. 2010; Wasan et al. 2009; Wasan et al. 2012; Weiss et al. 2014). However, epidemiological data indicate increased risk for opioid addiction in patients with chronic pain and increased prevalence of chronic pain in opioid users, including ~60% of opioid users or people with opioid addiction that experience chronic pain. (Cicero et al. 2008; Dahlhamer et al. 2018; Hser et al. 2017; John and Wu 2020; Juurlink and Dhalla 2012; Kaye et al. 2017a; Kaye et al. 2017b; Rosenblum et al. 2003).

In the present study, we found that across different experimental procedures, opioid drugs, pain-related manipulations, and labs, we consistently found a lack of effects of pain-related manipulations on opioid self-administration, reinstatement, and choice. Additionally, because females may be more sensitive to pain (Mogil and Bailey 2010), our intention was to follow up on positive findings with new experiments powered to detect sex differences, but we did not do this in light of our negative findings. Our results expand the range of preclinical conditions across which pain-related manipulations fail to increase opioid taking and seeking. These results suggest that increased prevalence of opioid addiction in patients with pain may be due to factors other than increased opioid reinforcement, such as increased availability of opioids through prescriptions or other means, decreased access to non-opioid rewards, co-morbid anxiety or depression, or pain-related cognitive deficits (Jensen et al. 2007; Kleiman et al. 2011; Moriarty et al. 2011; Nazarian et al. 2021). Recent efforts to better understand drug use patterns in humans (Panlilio et al. 2020) may reveal the importance of these factors for opioid use in patients with pain. This information could then be used to develop animal models to understand the complex relationship between pain and opioid use.

## Acknowledgements

We thank Hannah Korah for technical assistance with i.v. surgeries.

