## Supplemental Material for "Lack of effect of different pain-related manipulations on opioid self-administration, reinstatement of opioid seeking, and opioid choice in rats"

2-11-2021

**SUPPLAMENTAL ONLINE MATERIAL**

**Table of content**

Supplemental Methods

Figures S1-S5

Supplementary Table 1 (statistical analysis)

**Supplemental Methods**

Subjects

For Experiment 1-2 (conducted at IRP/NIDA/NIH, Baltimore, MD), we used 30 male and 18 female Sprague-Dawley rats (body weight at the time of arrival: males, 230-400 g; females, 200-240 g; Charles River). For Experiment 1A & 2A (food self-administration), we housed rats two per cage, and for Experiment 1B & 2B (opioid self-administration), we housed rats individually after i.v. surgery. We maintained the rats under a reverse 12:12 h light/dark cycle (lights off at 8 A.M.) with food (Teklad Rodent Diet, Envigo) and water freely available. In Experiment 1B, we excluded one male rat for poor health, and in Experiment 2B we excluded 4 rats for poor health (1 male, 3 females) and 5 female rats for catheter failure.

For Experiment 3 (conducted at Virginia Commonwealth University, Richmond, VA), we used 3 male and 5 female Sprague-Dawley rats (body weight at the time of arrival: 290-310g for males, 240-260g for females, Envigo). Following i.v. surgery, we housed the rats individually and maintained them on a 12-h light/dark cycle (lights off at 6:00 PM) with food (Teklad Rat Diet, Envigo) and water freely available. We performed the experiments in accordance with the NIH Guide for the Care and Use of Laboratory Animals (8th edition), under protocols approved by the NIDA IRP Animal Care and Use Committee or the Virginia Commonwealth University Institutional Animal Care and Use Committee.

Drugs

For Experiment 1-2, we obtained fentanyl citrate (fentanyl) and heroin hydrochloride (heroin) from the NIDA pharmacy and dissolved it in sterile saline. We chose a unit dose of 2.5 µg/kg/infusion for fentanyl self-administration training based on our previous study (Reiner et al. 2020). We chose unit doses of 0.1 and 0.05 mg/kg/infusion for heroin self-administration training based on previous studies (Bossert et al. 2016; Bossert and Stern 2014). We obtained capsaicin from Sigma-Aldrich (Cat#360376) as a powder, dissolved it with 10% ethanol and 10% Tween 80 in sterile saline, and injected it transdermally into the intraplantar region of the hindpaw in a volume of 50 µl. We obtained lactic acid from Sigma-Aldrich (Cat#252476), diluted it in sterile water to concentrations ranging from 0.9-1.8%, and injected it intraperitoneally (i.p.) in a volume of 1 ml/kg.

For Experiment. 3, we obtained fentanyl hydrochloride from NIDA Drug Supply Program (Bethesda, MD) and dissolved it in sterile saline. We obtained methohexital sodium from the Virginia Commonwealth University pharmacy, which we diluted in sterile water to a concentration of 16 mg/ml. We passed all i.v. solutions through a 0.22-micron sterile filter (Millex GV, Millipore Sigma) before administration. We expressed all drug doses as the salt forms listed above and delivered based on body weights collected weekly. We obtained lactic acid syrup (Cat#L1250) and Complete Freund’s Adjuvant (CFA, Cat#F5881) from Sigma Aldrich. We diluted lactic acid in sterile water to a 1% concentration and injected it (i.p.) in a volume of 1 ml/kg. We injected CFA transdermally into the intraplantar region the left hind paw in a volume of 0.1 ml.

Intravenous surgery

For Experiment 1-2, we anesthetized the rats with isoflurane gas (5% induction; 2-3% maintenance, Butler Schein) and inserted silastic catheters into the jugular vein, as previously described (Venniro et al. 2017a; Venniro et al. 2017b). We injected the rats with ketoprofen (2.5 mg/kg, s.c., Butler Schein) after surgery and the following day to relieve pain and inflammation. We allowed the rats to recover for 5-7 days prior to the experiment. During recovery and all experimental phases, we flushed the catheters every 24-48-h with gentamicin (4.25 mg/ml; APP Pharmaceuticals) dissolved in sterile saline.

For Experiment 3, we anesthetized rats with 2-3% isoflurane in oxygen and implanted them with polyurethane catheters into the right jugular vein using methods similar to those described previously (Townsend et al. 2015). We injected the rats with ketoprofen (5 mg/kg, s.c.) once immediately following surgery and again 24-h post-operatively. We allowed the rats to recover for 5 days prior to the experiment. After each behavioral session, we flushed catheters with 0.1 ml gentamicin (4 mg/ml; Aspen Veterinary Resources, Liberty, MO), followed by 0.1 ml of heparinized saline (10 U/ml).

Self-administration apparatus

For Experiment 1-2, we trained rats to self-administer food, fentanyl, or heroin in standard Med Associates (St. Albans, VT) self-administration chambers as described previously (Caprioli et al. 2015; Reiner et al. 2020). We equipped each self-administration chamber with two operant panels with three levers located 7-8 cm above the stainless-steel grid floor. We equipped the right panel of the chamber with a discriminative cue that signaled the insertion and subsequent availability of the food-paired active (retractable) lever. We equipped the left panel of the chamber with a discriminative cue that signaled the insertion and subsequent availability of the drug-paired active (retractable) lever. We also equipped the right wall with an inactive (stationary) lever that had no reinforced consequences.

For Experiment 3, we used 8 modular operant chambers located in sound-attenuating cubicles (Med Associates) equipped with two retractable levers, a set of three LED lights (red, yellow, green) mounted above each lever, and a retractable “dipper” cup (0.1 ml) located between the levers for presenting diluted Ensure® (32% v/v vanilla flavor Ensure® in tap water; Abbott Laboratories). Activation of a syringe pump delivered fentanyl solutions i.v. as described previously (Townsend et al. 2019b).

**General procedures**

Experiments 1-2

Food pellet self-administration: Prior to the first self-administration training session, we gave the rats 1-h magazine training session in which 1 pellet was delivered non-contingently every 3 min. We used 45-mg ‘preferred’ or palatable food pellets described in our previous studies (TestDiet, 1811155, 12.7% fat, 66.7% carbohydrate, and 20.6% protein) (Calu et al. 2014; Cifani et al. 2012; Pickens et al. 2012). We then trained rats to lever press for food during 1-h (Experiment 1A & 2B) or 3-h (Experiment 1B & 2B) sessions until they demonstrated reliable self-administration. The sessions began with the presentation of the white houselight, followed 10 s later by the insertion of the food-paired active lever. The white houselight remained on for the duration of the session and served as a discriminative cue for the palatable food. We trained the rats under a fixed-ratio-1 (FR1) 20-s timeout reinforcement schedule, where one lever press resulted in the delivery of one 45-mg palatable food pellet and the presentation of a 20-s discrete tone cue, during which additional lever presses were not reinforced but still recorded.

Drug self-administration: We trained rats to self-administer fentanyl or heroin under an FR1 20-s timeout reinforcement schedule (except where noted), where one lever press resulted in the delivery of a drug infusion paired with the 20-s discrete light cue above the drug-paired active lever. Sessions began with presentation of the houselight for 10 s followed by the insertion of the drug-paired active lever; the houselight remained on for the duration of the session and served as a discriminative cue for drug availability. At the end of each session, the houselight was turned off and the active lever was retracted.

Experiment 3

Choice experiments: We trained rats to respond in a fentanyl vs. food choice procedure as described previously (Townsend et al. 2019a; Townsend et al. 2019b). Briefly, we first trained rats to respond on the right lever for fentanyl (3.2 μg/kg/infusion) beginning under an FR1 20-s timeout schedule of reinforcement and progressing to an FR5 20-s timeout schedule of reinforcement. Illumination of a green stimulus light signaled fentanyl availability. Next, we similarly trained rats to respond on the left lever for a 5-s presentation of 32% Ensure® under an FR5 20-s timeout schedule of reinforcement. Illumination of a red stimulus light signaled Ensure® availability. Once rats responded for fentanyl and 32% Ensure® in isolation, we made both reinforcers available under a concurrent FR5 20-s timeout: FR5 20-s timeout schedule of reinforcement.

The behavioral session consisted of five 10-min response components, each preceded by a 4-min “sample” component. Each sample component started with a non-contingent infusion of the unit fentanyl dose available during the subsequent response component, followed by a 2-min timeout, and subsequently a 5-s presentation of liquid food, followed by a 2-min timeout. The response component began after this second timeout. During each response component, both levers extended, a red stimulus light above the left lever was illuminated to signal liquid food availability and a green stimulus light above the right lever was illuminated to signal iv fentanyl availability. Response requirement (FR5) completion on the left lever resulted in a 5-s presentation of liquid food whereas response requirement (FR5) completion on the right lever resulted in the delivery of the unit fentanyl dose available for that component. Responding on one lever reset the ratio requirement for the other lever.

We held the Ensure® concentration constant throughout the session, but varied the fentanyl dose during each of the five successive response components (0, 0.32, 1.0, 3.2, and 10 μg/kg/infusion during components 1–5, respectively) by changing the infusion duration (e.g., 315 g rat: 0, 0.5, 1.56, 5, and 15.6 s during components 1–5, respectively). To indicate a new fentanyl unit dose, the green light above the fentanyl-lever flashed on and off in 3-s cycles (i.e., longer flashes corresponded with larger fentanyl doses). During each response component, rats could complete up to 10 total ratio requirements between the food- and fentanyl-associated levers. Each ratio requirement completion initiated a 20 s time out, the retraction of both levers, and darkening of the red and green stimulus lights. If a rat completed all 10 ratio requirements before 10-min had elapsed, then both levers retracted, and stimulus lights were extinguished for the remainder of that component. We considered choice behavior stable when the smallest fentanyl dose that maintained at least 80% of completed ratio requirements on the fentanyl-associated lever was within a 0.5 log unit of the running mean for three consecutive days with no trends. We conducted fentanyl vs. food choice sessions five days per week from approximately 2:00 PM – 3:10 PM unless otherwise noted.

Mechanical sensitivity testing: We performed mechanical sensitivity testing approximately 3 h before the fentanyl-vs-food choice session for that day. We measured the width of the CFA-injected paw with calipers and then placed rats on an elevated mesh platform in individual chambers with a hinged lid. Following at least 20 min of acclimation, we exposed the rats to von Frey filaments (ranging from 0.4 to 15.0g and increasing in ~0.25 log increments; North Coast Medical, Morgan Hill, CA) on the plantar surface of each paw. We determined the threshold stimulus that elicited paw withdrawal in log grams using the “up-down” method as previously described (Chaplan et al. 1994; Leitl et al. 2014). We averaged threshold data for the injected paw across rats and thresholds for the noninjected paw almost always exceeded the 15 g ceiling (data not shown).

**Specific experiments**

**Experiment 1: Effect of intraplantar capsaicin on reinstatement of fentanyl seeking**

Experiment 1A: Effect of intraplantar capsaicin on food self-administration

We first trained 12 male rats to self-administer palatable food pellets for 1-h/day for 8 sessions. After the rats achieved stable food responding, we tested the effect of capsaicin on food self-administration. We anesthetized rats with isoflurane gas (5%) and injected 50 µg/50µl capsaicin or vehicle into the right hindpaw every other day based on previous studies demonstrating intraplantar injection of capsaicin increased mechanical and thermal hypersensitivity (Gilchrist et al. 1996; Hohmann et al. 2005). We immediately placed rats in the self-administration chamber and started a food self-administration session 30 min later to allow for adequate isoflurane recovery. To limit isoflurane exposure, we injected rats every other day of food self-administration for a total of 2 days of isoflurane exposure and food self-administration. We used an experimental design that included the between-subjects factor of capsaicin dose (0, 50 µg/50µl), n=6 per group and the within-subjects factor of Injection (first and second injection). We then used the six rats exposed to capsaicin to determine a capsaicin dose-response curve for depression of food self-administration in an experimental design that included the within-subjects factor of capsaicin dose (0, 50, or 100 µg/50 µl).

Experiment 1B: Effect of intraplantar capsaicin context on fentanyl seeking

Fentanyl self-administration: Rats from Experiment 1A were used in Experiment 1B. We performed i.v. surgery on 11 of the male rats and used a different behavioral room and self-administration chambers. We trained the rats to self-administer fentanyl for 12 days in two 1-h daily sessions, separated by a 10-min timeout period. Fentanyl was infused at a dose of 2.5 µg/kg/infusion over 3.5 s (0.1 ml/infusion) followed by a 20-s timeout period. This fentanyl unit dose is on the peak or descending limb of fentanyl self-administration dose-effect curves (Martin et al. 2007; Townsend et al. 2019b; Wade et al. 2015). We trained rats to self-administer in either No Capsaicin or Capsaicin contexts, which differed in color of the discriminative houselight (white or red), color of the fentanyl-paired cue light (white or red), thickness of the grid floor, type of palatable food pellet dispenser, and presence of empty water bottle and food hopper. We counterbalanced the physical environments of No Capsaicin and Capsaicin contexts and alternated No-Capsaicin and Capsaicin contexts every other day (counterbalanced across rats), for a total of 6 self-administration days in each context. For the Capsaicin context, we anesthetized rats with isoflurane gas (5%) and injected 100 µg/50µl capsaicin in the right hindpaw. We immediately placed the rats in the self-administration chamber and started the fentanyl self-administration session 30 min later to allow for adequate isoflurane recovery. We did not expose rats in the No-Capsaicin context to isoflurane and started the fentanyl self-administration session after 30 min of habituation in the self-administration chamber.

Capsaicin dose response: Next, we determined a capsaicin dose-response curve on fentanyl self-administration by injecting capsaicin in the Capsaicin context for four consecutive days. We used a counterbalanced experimental design that included the within-subjects factor of Capsaicin dose (0, 50, 100, or 200 µg/50 µl).

Extinction: Next, we exposed the rats to extinction conditions in which responses on the previously active lever led to presentation of the fentanyl-paired cue light, but fentanyl was not delivered. We also alternated No-Capsaicin and Capsaicin contexts (counterbalanced), for a total of 8 extinction days, 4 in each context. In the previous Capsaicin context, we exposed the rats to ~1 min of 5% isoflurane exposure prior to beginning the extinction session. For both contexts, we started each session 30 min after placing rats in the chamber.

Reinstatement: We tested rats under extinction conditions (active lever presses led to the presentation of the cue light but no fentanyl infusions) for two 1-h sessions over 3 days in a counterbalanced order. All rats (n=11) were exposed to the No-Capsaicin context, 6 of the 11 rats were exposed to vehicle in the Capsaicin context, and 5 of the 11 rats were exposed to capsaicin (100 µg/50 µl) in the Capsaicin context. We did not reverse the conditions in the Capsaicin context because the initial test indicated that capsaicin injections had no effect on reinstatement.

**Experiment 2: Effect of i.p. lactic acid on reinstatement of heroin seeking**

Experiment 2A: Effect of i.p. lactic acid on food self-administration

FR1 food self-administration: We determined the effect of i.p. lactic acid injections on food self-administration. We trained 4 male and 4 female rats to self-administer palatable food during daily 3-h sessions for 5 days, habituating them to vehicle (sterile water, i.p.) injections before the last 2 sessions. For all lactic acid experiments, we injected rats with vehicle or lactic acid (i.p.) and started the session 5 min later. We injected 1.8% lactic acid every other day of self-administration for a total of 3 days of 1.8% injections and 2 no injection days, followed by 1 day of vehicle injections. We used an experimental design that included the within-subject factor of Lactic acid concentration (0, 1.8%). To examine the effect of 0.9% i.p. lactic acid on food self-administration, we then injected 0.9% lactic acid every other day of self-administration for a total of 3 days of 0.9% injections and 2 no injection days, followed by 1 day of vehicle injections. We used an experimental design that included the within-subject factor of Lactic acid concentration (0, 0.9%). We based these doses on previous studies demonstrating lactic acid depressed intracranial self-stimulation and increased nociceptive behaviors such as stretching (Pereira Do Carmo et al. 2009).

Progressive ratio (PR) self-administration: We next tested the effect of i.p. lactic acid on food self-administration using a PR schedule of reinforcement. We trained rats on this PR schedule for 3 days, habituating them to vehicle injections on the last day. We injected i.p. lactic acid every other day of self-administration for a total of 3 lactic acid injections (0.9, 1.35, 1.8%) and 2 vehicle injection days. We used an experimental design that included the within-subject factor of lactic acid concentration (0, 0.9, 1.35, 1.8%). The PR ratio increments were: 2, 4, 6, 9, 12, 15, 20, 25, 32, 40, 50, 62, 77, 95, 118, 145, 178, 219, and so on (Richardson and Roberts 1996).

Ethogram: During PR self-administration testing, we also measured the effect of i.p. lactic acid on pain behaviors, which we defined as dragging (dragging lower half on ground for short increments), immobile (standing without locomotion or apparent sniffing, grooming, or other movement), laying (curled up, lying on side), pancaking (lying on floor, stretched out so that abdomen touches the floor), stretching (stretching/writhing), licking abdomen (licking injection site), hunched posture (standing with paws close to each other and back bone arched and lifted), and no-pain behaviors, which we defined as scratching (any limb), grooming (licking fur or washing face with forepaws), rearing (on hind legs), sniffing, locomotion, still and alert (sitting but face is lifted) (Roughan and Flecknell 2003). We counted the number of behaviors per min and summed these behaviors to determine the number of No pain- and Pain-related behaviors for each rat within the first 30 min. We used an experimental design that included the within-subject factors of lactic acid concentration (0, 0.9, 1.35, 1.8%) and behavior type (No pain, Pain).

Experiment 2B: Effect of i.p. lactic acid on reinstatement of heroin seeking

Self-administration and extinction: We trained rats (n=13 males and 6 females) to self-administer palatable food pellets for 3-h/day for 4 days, followed by heroin self-administration for 3-h/day for 19 days. Heroin was infused at a volume of 100 µl over 3.5 s at a dose of 0.1 mg/kg/infusion (the first 11 sessions) and then 0.05 mg/kg/infusion (last 9 sessions). These heroin unit doses are on the descending limb of the heroin self-administration dose-effect curve (Martin et al. 2007; Stewart et al. 1996). We trained rats to self-administer heroin paired with a light cue above the active lever (red lens) under an FR1 20-s timeout reinforcement schedule (the first 12 sessions) and then increased the response requirement to FR3 for 2 sessions, FR6 for 2 sessions, and returned the response requirement to FR3 for the last 3 sessions. The rats received an i.p. injection of either vehicle (sterile water, n=7) or 0.9% lactic acid (n=13) 5 min prior to the start of each heroin self-administration session. We then extinguished responding for heroin for 12 days without i.p. vehicle or lactic acid exposure; during the extinction sessions, responses on the previously active lever led to presentations of the heroin-paired cue light, but not heroin infusions. We habituated the rats to priming saline injections (s.c.) during the last day of extinction before heroin priming. We included self-administration data from one rat in the vehicle training condition but eliminated this rat from subsequent phases of the experiment because of poor health during the extinction phase.

Effect of lactic acid on reinstatement of heroin seeking: We tested the effect of i.p. lactic acid on reinstatement of heroin seeking during three 3-h sessions under extinction conditions in the two groups of rats previously exposed to vehicle (n=6) or lactic acid (n=13) during training. We first injected the vehicle (i.p.) and two concentrations of i.p. lactic acid (0.45 and 0.9%) in a counterbalanced order. After these 3 sessions, we also tested the effect of 1.35% lactic acid on heroin seeking.

Effect of lactic acid on heroin priming-induced reinstatement of heroin seeking: For the rats exposed to i.p. lactic acid during training (n=13), we used a between-subjects design in which the rats were pretreated with either i.p. vehicle (n=6) or i.p lactic acid (0.9%, n=7) immediately before heroin priming injections (0.25 mg/kg, s.c) 10 min before the start of the session. For the rats exposed to i.p. vehicle during training (n=6), we used a within-subjects design in which the rats were pretreated with i.p. vehicle and lactic acid (0.9%) before the heroin priming injections. We switched to using a within-subjects design, because of a disproportional loss of subjects in the vehicle training group. We based the heroin priming dose on prior studies on reinstatement of heroin seeking (Shaham et al. 1996; Shaham and Stewart 1995) and a pilot study which showed that this dose led to reliable and robust reinstatement.

**Experiment 3: Effect of i.p. lactic acid and intraplantar CFA on choice fentanyl vs. food choice**

Experiment 3A: Effect of repeated i.p. lactic acid injections on fentanyl vs. food choice

We trained 3 male and 5 female rats on the fentanyl vs. food choice procedure. Once fentanyl vs. food choice behavior stabilized, we injected either 1 ml/kg of 1% i.p. lactic acid or volume-matched vehicle (sterile water) injections immediately prior to fentanyl vs. food choice tests for five consecutive sessions (i.e., 5 consecutive i.p. lactic acid injections or 5 consecutive vehicle injections). We counterbalanced the order of lactic acid and vehicle testing across rats. We based the i.p. lactic acid concentration and repeated treatment regimen on previous studies showing that 1% i.p. lactic acid is the approximate threshold for producing repeatable pain-related and opioid-reversible decreases in positively reinforced operant responding in rats (Miller et al. 2015; Pereira Do Carmo et al. 2009).

Experiment 3B: Effect of intraplantar CFA on fentanyl vs. food choice, mechanical sensitivity, and paw width

During the final two days (Thursday & Friday) of the repeated i.p. lactic acid and vehicle experiment, we assessed mechanical sensitivity and paw width 3-h prior to the daily fentanyl vs. food choice tests to acclimate rats to the procedure and serve as the “baseline” for subsequent analyses of mechanical sensitivity and paw width. We performed a “baseline” fentanyl vs. food choice session on Sunday, and then on the next day briefly sedated the rats with 3% isoflurane in oxygen and injected the rats intradermally with 0.1 ml of intraplantar CFA into the left hindpaw. We then performed fentanyl vs. food choice tests Monday through Friday for three weeks (i.e., up to 18 days post-CFA) and on Mondays only for the next two weeks (i.e., 21 and 28 days post-CFA). We assessed mechanical sensitivity and paw width 1, 3, 7, 14, 21, and 28 days post-CFA at least one hour before the daily fentanyl vs. food choice session. We chose the volume of intraplantar CFA because this volume of CFA produces sustained mechanical hypersensitivity and transient depression of positively reinforced operant responding in rats (Leitl et al. 2014).

Experiment 3C: Effect of FR manipulations on fentanyl vs. food choice

After day 28 post-CFA treatment, we independently increased the response requirement for food and fentanyl injections from 5 to 30 for two consecutive sessions in a counterbalanced order. We used the second day of each response requirement manipulation for subsequent analyses. We verified catheter patency in all rats at the conclusion of this final experiment by instantaneous loss of muscle tone following IV methohexital (1.6 mg) administration.

Statistical analysis

In Experiments 1-2, we analyzed the data with repeated-measures or mixed-factorial ANOVAs using SPSS (Version 23, GLM procedure). We followed significant main effects and interactions (p<0.05) with post-hoc tests (univariate ANOVAs or Fisher’s PLSD). We describe the different between- and within-subjects factors for the different statistical analyses in the Results section. Unless otherwise indicated, we included some analyses using average number of rewards or average number of lever presses after determining no significant differences between sessions. We only report significant effects that are critical for data interpretation, as our multifactorial ANOVAs yielded multiple main and interaction effects. We indicate the results of post-hoc analyses with asterisks in the figures, but do not describe them in the text. For a complete reporting of the statistical analyses see supplemental Table S1.

In Experiment 3, we analyzed four primary dependent measures from the fentanyl choice self-administration sessions: (1) “percent component fentanyl choice” defined as [(number of ratio requirements, or ‘choices’, completed on the fentanyl-associated lever ÷ total number of choices completed on both the drug- and food-associated levers during each component) × 100], (2) “reinforcement rate per component” defined as total number of choices completed during each component, (3) percent session fentanyl choice, defined as [(number of fentanyl choices completed ÷ total number of drug and food choices completed for the entire session) × 100], and (4) total, food, and fentanyl choices completed during the entire 2-h session. Parameters of fentanyl choice, as well as measures of mechanical sensitivity and paw width, were analyzed using one-way or two-way repeated-measures ANOVA with drug dose, experimental manipulation, or time as the main factors and used the Geisser-Greenhouse correction when appropriate (Prism 8, GraphPad). We followed up on significant main effects or interactions with post-hoc tests appropriate for the pre-planned comparisons and corrected for multiple comparisons.

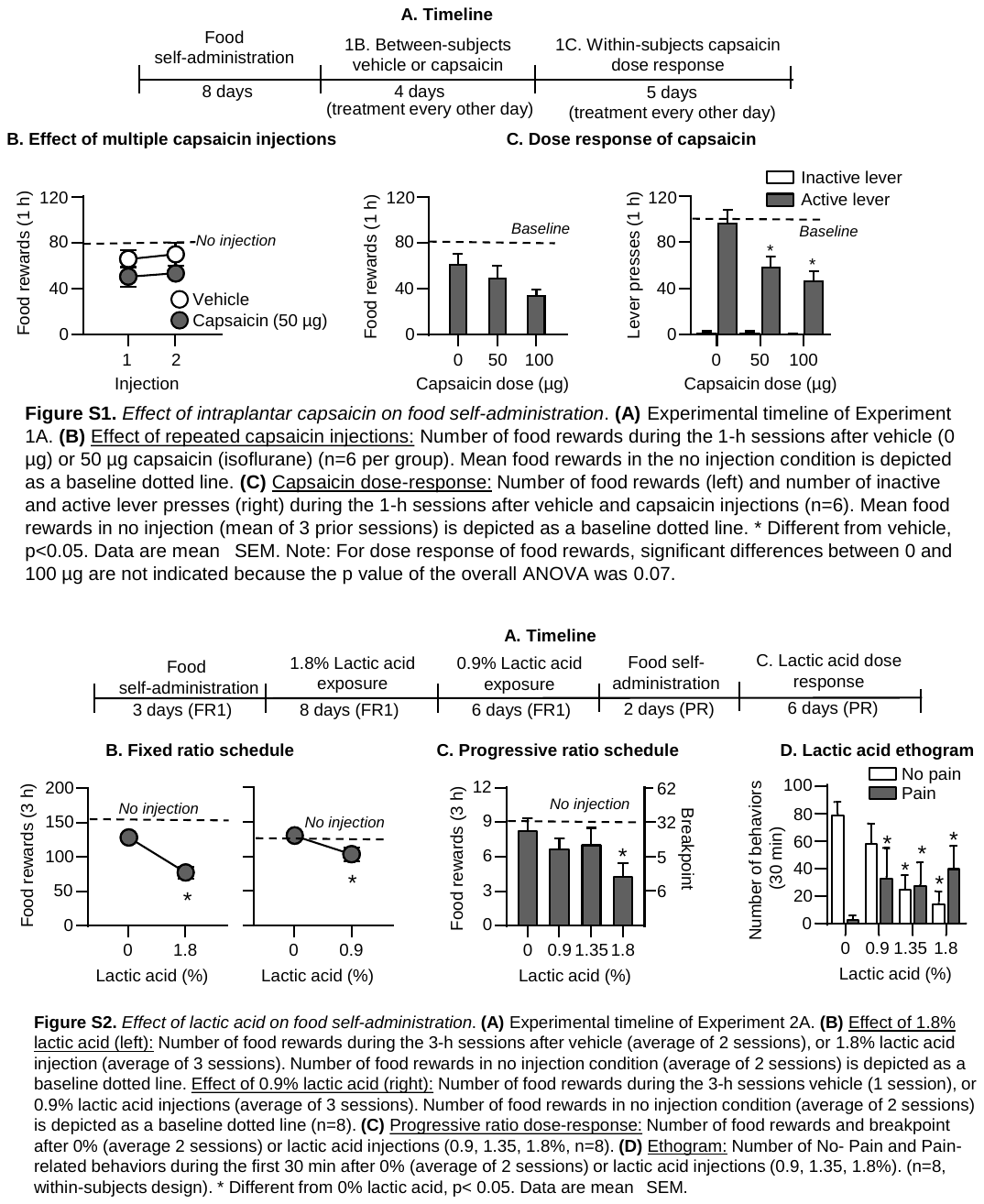

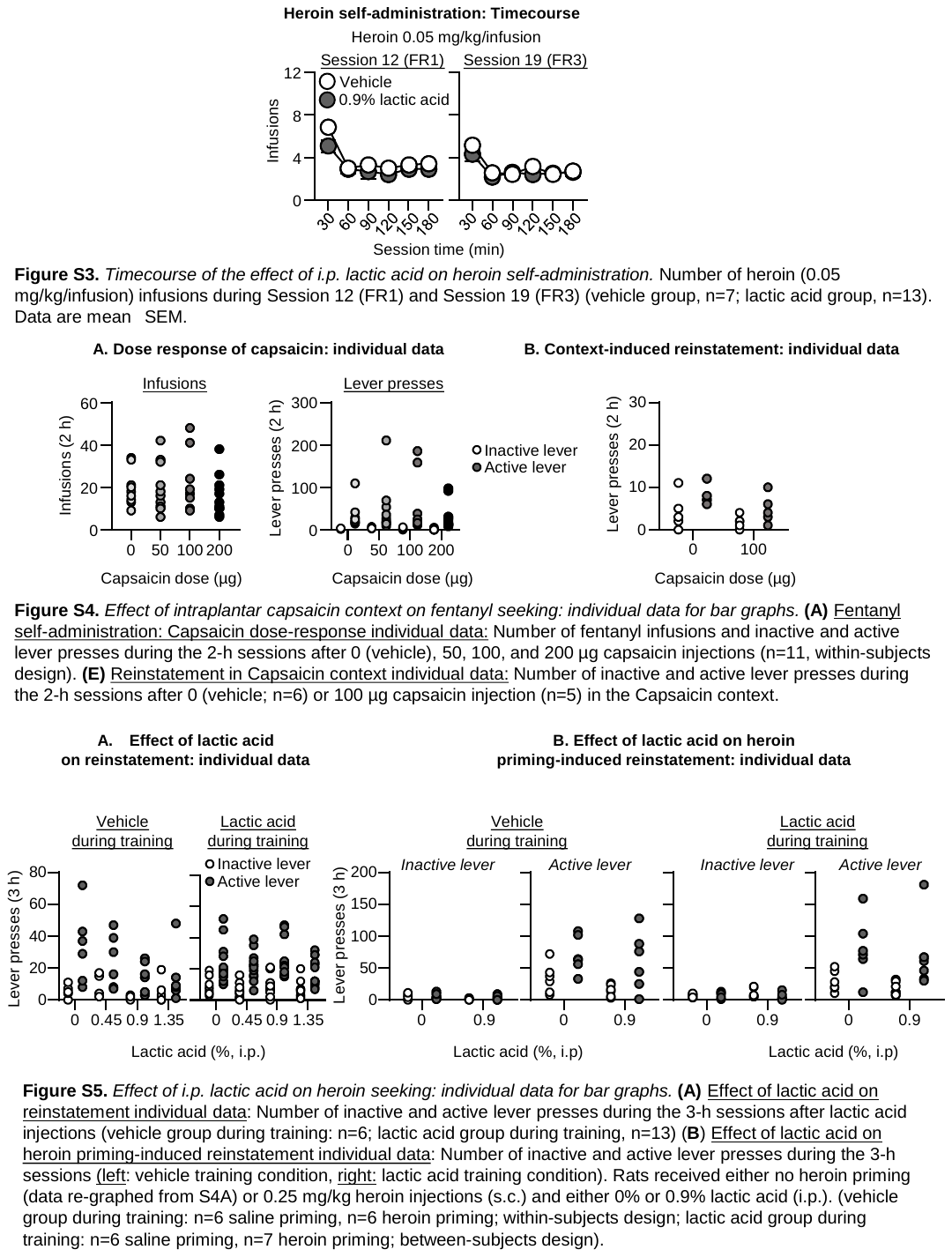

**Supplementary Table 1.** Statistical analysis for Experiments 1-2 (SPSS GLM repeated-measures module) and Experiment 3 (Prism 8, GraphPad). Partial Eta^2^ = proportion of explained variance. NP = not possible.

| **Figure number** | **Factor name** | **F-value** | ***p*-value** | **Partial Eta^2^** |
| --- | --- | --- | --- | --- |
| Figure 1B. Fentanyl self-administration: No pain and Pain context | **Infusions**  Context (No pain, Capsaicin), within-subjects  Session (1-6), within-subjects  Context x Session  **Lever presses**  Context (No pain, Capsaicin), within-subjects  Session (1-6), within-subjects  Lever (Inactive, active), within-subjects  Context x Session  Context x Lever  Session x Lever  Session x Context x Lever | F_1,10_=0.2  F_5,50_=5.6  F_5,50_=1.0  F_1,10_=0.4  F_5,50_=1.3  F_1,10_=14.3  F_5,50_=0.5  F_1,10_=0.2  F_5,50_=3.3  F_5,50_=0.5 | 0.61  <0.001*  0.37  0.54  0.28  0.004*  0.78  0.67  0.01*  0.78 | 0.02  0.35  0.09  0.04  0.12  0.59  0.05  0.02  0.25  0.05 |
| Figure 1C. Fentanyl self-administration: Capsaicin dose response | **Infusions**  Capsaicin dose (0, 50, 100, or 200 µg), within-subjects  **Lever presses**  Capsaicin dose (within-subjects)  Lever (Inactive, active), within-subjects  Capsaicin dose x Lever | F_3,30_=1.7  F_3,30_=1.2  F_1,10_=8.2  F_3,30_=1.2 | 0.18  0.32  0.02*  0.33 | 0.14  0.11  0.45  0.11 |
| Figure 1D. Extinction in No pain and Pain context | **Lever presses**  Session (1-4), within-subjects  Context (No pain, Pain), within-subjects  Lever (Inactive, active), within-subjects  Session x Lever  Context x Lever  Session x Context x Lever | F_3,30_=15.0  F_1,10_=0.0  F_1,10_=26.8  F_3,30_=15.5  F_1,10_=0.0  F_3,30_=0.5 | <0.001*  0.88  <0.001*  <0.001*  0.96  0.68 | 0.60  0.002  0.72  0.60  0.000  0.04 |
| Figure 1E. Reinstatement in No pain and Pain context | **Lever presses**  Capsaicin dose (0, 100 µg), between-subjects  Lever (Inactive, active), within-subjects  Interaction | F_1,9_=2.3  F_1,9_=9.4  F_1,9_=1.4 | 0.16  0.013*  0.67 | 0.20  0.51  0.022 |
| Figure 2B. Heroin Self-administration | **Infusions**  Training condition (vehicle, 0.9% lactic acid), between-subjects  Session (Sessions 1-19), within-subjects  Interaction | F_1,18_=0.7  F_18,324_=12.9  F_18,324_=0.6 | 0.41  <0.001*  0.92 | 0.04  0.41  0.03 |
| Figure 2C. Heroin Extinction | **Lever presses**  Training condition (vehicle, 0.9% lactic acid), between-subjects  Lever (active, inactive), within-subjects  Session (1-12), within-subjects  Training condition x Lever  Lever x Session  Training condition x Session  Training condition x Lever x Session | F_1,17_=0.05  F_1,17_=104.5  F_11,187_=32.1  F_1,17_=0.26  F_11,187_=25.7  F_11,187_=0.68  F_11,187_=0.46 | 0.83  <0.001*  <0.001*  0.62  <0.001*  0.75  0.93 | 0.00  0.86  0.65  0.02  0.60  0.04  0.03 |
| Figure 2D. Effect of lactic acid on reinstatement Relapse test | **Lever presses**  Training condition (vehicle, 0.9% lactic acid), between-subjects  Lactic acid conc. (0.0, 0.45, 0.9, 1.35%), within-subjects  Lever (Inactive, active), within-subjects  Training condition x Lactic acid conc.  Training condition x Lever  Lactic acid conc. x Lever  Lactic acid conc. x Training condition x Lever  Post hoc: vehicle training group:  Lactic acid conc., within-subjects  Lever, within-subjects  Lactic acid conc. x Lever  Post hoc: lactic acid training group:  Lactic acid conc., within-subjects  Lever, within-subjects  Lactic acid conc. x Lever | F_1,17_=0.2  F_3,51_=4.9  F_1,17_=43.3  F_3,51_=4.5  F_1,17_=0.11  F_3,51_=5.4  F_3,51_=2.9  F_3,15_=3.0  F_1,5_=13.3  F_3,15_=3.7  F_3,36_=4.2  F_1,12_=35.6  F_3,36_=1.7 | 0.66  0.004*  <0.001*  0.007*  0.75  0.003*  0.041*  0.06  0.02*  0.035*  0.012*  <0.001*  0.18 | 0.01  0.23  0.72  0.21  0.01  0.24  0.15  0.38  0.73  0.43  0.26  0.75  0.13 |
| Figure 2E. Effect of lactic acid on heroin priming-induced reinstatement test | **Active lever presses (Vehicle training group)**  Heroin priming (no priming, 0.25 mg/kg), within-subjects  Lactic acid (0, 0.9%), within-subjects  Lever (Inactive, active), within-subjects  Heroin priming x Lactic acid  Heroin priming x Lever  Lever x Lactic acid  Heroin priming x Lever x Lactic acid | F_1,5_=8.9  F_1,5_=3.6  F_1,5_=36.4  F_1,5_=0.2  F_1,5_=7.4  F_1,5_=1.54  F_1,5_=0.07 | 0.03*  0.12  0.002*  0.67  0.04*  0.27  0.80 | 0.64  0.42  0.88  0.04  0.60  0.24  0.01 |
| Figure 2E. Effect of lactic acid on heroin priming-induced reinstatement test | **Active lever presses (Lactic acid training group)**  Heroin priming (no priming, 0.25 mg/kg), within-subjects  Lactic acid (0, 0.9%), between-subjects  Lever (Inactive, active), within-subjects  Heroin priming x Lactic acid  Heroin priming x Lever  Lever x Lactic acid  Heroin priming x Lever x Lactic acid | F_1,11_=12.0  F_1,11_=0.5  F_1,11_=30.2  F_1,11_=0.04  F_1,11_=17.4  F_1,11_=0.66  F_1,11_=0.04 | 0.005*  0.49  <0.001*  0.85  0.002*  0.43  0.85 | 0.52  0.04  0.73  0.00  0.61  0.06  0.00 |
| Figure 3B (left). Effect of lactic acid on fentanyl-vs.-food dose response | **Percent fentanyl choice dose response**  Unit dose (0, 0.32, 1, 3.2, 10), within subjects  Lactic acid concentration (0, 1%), within subjects  Interaction | F_1.6, 11.1_=50.7  F_1.0, 7.0_=0.75  F_2.3, 12.7_=0.50 | <0.0001*  0.42  0.64 | NP  NP  NP |
| Figure 3B (right). Effect of lactic acid on fentanyl-vs.-food across sessions | **Fentanyl choice across sessions**  Time (sessions 1-5), within subjects  Lactic acid concentration (0, 1%), within subjects  Interaction | F_2.3, 16.4_=0.72  F_1.0, 7.0_=3.8  F_2.8, 18.3_=0.91 | 0.52  0.09  0.45 | NP  NP  NP |
| Figure 3C (left). Effect of lactic acid on reinforcement rate dose response | **Reinforcement rate dose response**  Unit dose (0, 0.32, 1, 3.2, 10), within subjects  Lactic acid concentration (0, 1%), within subjects  Interaction | F_2.1, 14.7_=51.3  F_1.0, 7.0_=16.9  F_1.8, 12.5_=9.0 | <0.0001*  0.0045*  0.0045* | 0.94  0.75  0.56 |
| Figure 3C (right). Effect of lactic acid on reinforcement rate across sessions | **Reinforcement rate across sessions**  Time (sessions 1-5), within subjects  Lactic acid concentration (0, 1%), within subjects  Interaction | F_2.2, 15.6_=1.0  F_1.0, 7.0_=17.2  F_2.2, 15.7_=0.55 | *p*=0.39  *p*=0.004*  *p*=0.61 | 0.12  0.55  0.07 |
| Figure 4B (left). Effect of CFA on fentanyl-vs.-food dose response | **Percent fentanyl choice dose response**  Fentanyl unit dose 0, 0.32, 1, 3.2, 10), within subjects  Time since CFA (-1, 1, 3, 7), within subjects  Interaction | F_1.6, 11.0_=36.8  F_0.84, 5.9_=0.64  F_2.2, 9.6_=1.4 | <0.0001*  0.42  0.30 | NP  NP  NP |
| Figure 4B (right). Effect of CFA on fentanyl-vs.-food across sessions | **Fentanyl choice across sessions**  Days since CFA (sessions -1,0-4, 7-11, 14), within subjects | F_3.6, 25.5_=1.1 | 0.37 | 0.14 |
| Figure 4C (left). Effect of CFA on reinforcement rate dose response | **Reinforcement rate dose response**  Fentanyl unit dose (0, 0.32, 1, 3.2, 10), within subjects  Time since CFA (-1, 1, 3, 7), within subjects  Interaction | F_1.9 ,13.0_=73.9  F_1.3, 8.9_=6.0  F_2.9, 20.1_=2.5 | <0.0001*  0.03*  0.09 | 0.79  0.38  0.26 |
| Figure 4C (right). Effect of CFA on reinforcement rate across sessions | **Reinforcement rate across sessions**  Days since CFA (sessions -1,0-4, 7-11, 14), within subjects | F_2.3, 16.0_=4.5 | 0.02* | 0.39 |
| Figure 4D (left). Effect of CFA on mechanical sensitivity. | **Mechanical sensitivity**  Days since CFA (sessions -1,1, 3, 7, 14), within subjects | F_2.1, 14.5_=17.5 | 0.0001* | 0.71 |
| Figure 4C (right). Effect of CFA on paw width | **Paw width**  Days since CFA (sessions -1,1, 3, 7, 14), within subjects | F_2.4, 16.7_=100.9 | 0.0001* | 0.94 |
| Figure 5B (left). Effect of FR manipulation on fentanyl-vs.-food dose response | **Percent fentanyl choice dose response**  Fentanyl unit dose (0, 0.32, 1, 3.2, 10), within subjects  FR (FR5:FR5; FR30:FR5; FR5; FR30) within subjects  Interaction | F_2.0, 14.2_=30.4  F_1.9, 13.3_=66.4  F_3.7, 22.1_=7.9 | <0.0001*  <0.0001*  0.0005* | NP  NP  NP |
| Figure 5B (right). Effect of FR manipulation on reinforcement rate dose response | **Reinforcement rate dose response**  Fentanyl unit dose (0, 0.32, 1, 3.2, 10), within subjects  FR (FR5:FR5; FR30:FR5; FR5; FR30) within subjects  Interaction | F_2.1, 14.9_=85.0  F_1.6, 11.0_=12.4  F_2.4, 16.9_=1.2 | <0.0001*  0.002*  0.35 | 0.87  0.70  0.66 |
| Figure S1B. Effect of capsaicin | **Rewards**  Injection (1, 2), within-subjects  Capsaicin dose (Vehicle, capsaicin), between-subjects  Sessions x Capsaicin dose | F_1,10_=0.4  F_1,10_=2.9  F_1,10_=0.01 | 0.53  0.12  0.91 | 0.04  0.22  0.001 |
| Figure S1C. Capsaicin dose response (within-subjects) | **Rewards**  Capsaicin dose (0, 50, or 100 µg), within-subjects  **Lever presses**  Capsaicin dose (0, 50, or 100 µg), within-subjects  Lever (Inactive, active), within-subjects  Interaction | F_2,10_=3.5  F_2,10_=9.5  F_1,5_=122.9  F_2,10_=10.3 | 0.07  0.005*  <0.001*  0.004* | 0.41  0.66  0.96  0.67 |
| Figure S2B. Effect of 1.8% lactic acid | **Rewards**  Lactic acid concentration (0, 1.8%), within-subjects | F_1,7_=89.5 | <0.001* | 0.92 |
| Figure S2B. Effect of 0.9% lactic acid | **Rewards**  Lactic acid concentration (0, 0.9%), within-subjects | F_1,7_=6.6 | 0.037* | 0.48 |
| Figure S2C. Progressive ratio self-administration | **Rewards**  Lactic acid concentration (0, 0.9, 1.35, 1.8%), within-subjects | F_3,21_=6.0 | 0.004* | 0.46 |
| Figure S2D. Ethogram | **Number of Behaviors:**  Behavior type (No pain, Pain), within-subjects  Lactic acid concentration (0, 0.9, 1.35, 1.8%), within-subjects  Interaction | F_1,7_=10.5  F_3,21_=46.6  F_3,21_=36.1 | 0.014*  <0.001*  <0.001* | 0.60  0.87  0.83 |
| Figure S3. Within-session heroin Self-administration  FR1  FR3 | **Infusions**  Training condition (0, 0.9% lactic acid), between-subjects  Session time (30,60,90,120,150,180), within-subjects  Training condition x Session time  Training condition, between-subjects  Session time, within-subjects  Training condition x Session time | F_1,18_=1.8  F_5,90_=20.9  F_5,90_=1.8  F_1,18_=0.4  F_5,90_=18.0  F_5,90_=1.1 | 0.19  <0.001*  0.11  0.52  <0.001*  0.38 | 0.09  0.54  0.09  0.02  0.5  0.06 |
